# Neuroanatomical Subtypes of Callous-Unemotional Traits in a Community Sample of Youth

**DOI:** 10.64898/2026.07.14.738524

**Authors:** K. Murtha, M. Antonaides, J. Seidlitz, R. Barzilay, T.M. Moore, R. Shinohara, T.D. Satterthwaite, E.R. Kimonis, C. Davatzikos, R. Waller

## Abstract

**Importance:** Callous Unemotional (CU) traits are associated with significant clinical and neurophysiological heterogeneity that may affect treatment effectiveness.

**Objective:** To uncover neuroanatomical subtypes of CU traits using weakly-supervised machine learning and assess whether emerging subtypes differ on relevant clinical, temperamental, and environmental constructs.

**Design, Setting, and Participants:** Imaging data was from the longitudinal Adolescent, Brain, Cognitive Development (ABCD) Study. Participants were 9-10 years old at baseline (M=9.925, 69.9% male) and included 222 children with CU traits and 234 typically developing controls matched on age, sex and income.

**Main Outcomes and Measures:** The weakly-supervised heterogeneity through discriminative analysis (HYDRA) model was trained on grey matter (GM) volumes from 84 regions of interest (ROIs) and tested for reproducibility using cross-validation and permutation testing. Derived subtypes were compared cross-sectionally and prospectively on relevant clinical, temperamental, and environmental measures and subsequent GM volume at 2-year follow up.

**Results:** HYDRA revealed an optimal 2-subtype solution within children with CU traits. Subtypes showed comparable levels of aggression that were significantly higher than typically developing controls. At the same time, subtype 1 had larger GM volume, fewer internalizing symptoms, and less adversity exposure, while subtype 2 was characterized by smaller GM volume, more internalizing symptoms, and more adversity exposure.

**Conclusions and Relevance:** This study provides evidence of neuroanatomically distinct subtypes of CU traits characterized by different clinical and etiological profiles, with implications for diagnosis and treatment.

## Introduction

Callous-unemotional (CU) traits are defined by low empathy, guilt, or prosociality^1,2^ and predict very high risk for childhood disruptive behavior disorders (DBDs) and antisocial behavior across the lifespan,^3^ creating a vast economic and health burden.^4–6^ Research has identified early adverse experiences,^7^ genetic vulnerabilities,^8^ and neurophysiological factors^9,10^ that play a role in the development and maintenance of CU traits. More specifically, CU traits are associated with structural abnormalities^11^ and disrupted functioning in the amygdala, particularly in relation to fearful stimuli,^12,13^ which aligns with their characteristic low sensitivity to threat.^14,15^ CU traits have also been related to volumetric differences in cortical regions, including differences in anterior temporal lobe, orbitofrontal, and anterior cingulate cortex gray matter volume (GMV), which may indicate delays in cortical maturation in brain regions related to decision making and empathy.^16,17^

However, the CU traits phenotype itself is heterogeneous, with a growing number of studies focused on distinct primary and secondary subtypes thought to arise through unique etiological pathways.^18,19^ This literature draws on early theories of psychopathy in adults describing a distinction between “idiopathic” (primary) psychopathy, which was hypothesized to arise from inherited neurophysiological differences, and “symptomatic” (secondary) psychopathy, which was believed to develop following experiences of abuse or neglect in early life.^20,21^ Likewise, primary CU traits are hypothesized to arise from genetically-mediated fearlessness and low neurophysiological arousal.^15^ In contrast, secondary CU traits are thought to emerge in the context of heightened neurophysiological arousal and greater exposure to interpersonal adversity and trauma (i.e., “reactionary callousness”) that produce complex co-occurring mental health challenges, including anxiety and trauma symptoms.^22–24^

Most commonly, primary and secondary subtypes of CU traits are identified based on having low versus high levels of co-occurring internalizing symtomatology.^18^ Some studies have also identified theory-consistent differences in the physiological correlates of primary and secondary CU traits subtypes, including differences in cortisol or startle response.^25,26^ However, a paucity of studies have examined neurobiological differences. Limited evidence suggests that children with secondary CU traits demonstrate lower functional connectivity between the amygdala and various cortical regions,^27^ while children with primary CU traits showed diminished amygdala response to neutral^28^ and fearful faces.^29^ No prior studies have tested brain structural differences between subtypes. However, children with secondary CU traits may exhibit similar reductions in cortical and subcortical volume to those reported by studies on GMV abnormalities among children exposed to adversity or with anxiety.^30–32^

Importantly, prior studies of primary and secondary subtypes have largely relied on parent or youth report measures to both *derive* and *validate* CU traits subtypes,^24,33^ which inflates shared method variance and raises concerns about circularity in validating subtypes, potentially undermining construct and external validity.^34,35^ As a result, existing evidence provides limited insight into whether these subtypes show distinct neurobiological profiles consistent with hypothesized primary versus secondary etiological pathways to CU traits.^36^ Weakly-supervised machine learning may offer a more effective method to uncover clinically meaningful subtypes by modeling patterns of deviation from a reference group (e.g., typically developing [TD] control), using a data-driven, bottom-up analytic approach.^37^ Although these methods have been effectively applied in other clinical populations, including Autism Spectrum Disorder and depression,^38–40^ they have not been used to study heterogeneity in disruptive behavior disorders or CU traits.

Thus, the current study aimed to leverage weakly-supervised machine learning to identify unique profiles of primary and secondary CU traits, which may better characterize children based on their underlying neurobiology. We hypothesized that (1) distinct profiles of cortical (cGMV) and subcortical (sGMV) grey matter volume would differentiate subgroups within children exhibiting symptoms of disruptive behavior disorders and CU traits, (2) that subtypes would differ on levels of internalizing (e.g., anxiety and depression) symptoms in line with theory driven definitions of primary and secondary CU traits, (3) a subtype with high levels of internalizing symptoms would also have experienced greater adversity; and (4) subtypes would evidence distinct temperamental profiles characterized by sensitivity to threat and affiliative reward.

## Methods

### Participants

Participants were a subsample of the Adolescent Brain Cognitive Development study, a large-scale longitudinal study of over 11,000 children across 21 sites in the United States with baseline data collection at ages 9-10 years old.^41^ Participants underwent structural magnetic resonance imaging (sMRI) alongside clinical phenotyping using parent and youth report. We used data from the baseline and longitudinal 2-year follow-up assessments.

### Deriving Clinical and Typically Developing Groups

We first used baseline diagnostic data from the Kiddie Schedule for Affective Disorders and Schizophrenia (K-SADS)^42^ to identify children whose parents reported one or more disruptive behavior disorders (DBD, e.g., conduct disorder, oppositional defiant disorder). Of those children, we used a validated 4-item measure of CU traits to identify 258 children who scored at or above the 95^th^ percentile (i.e., scores ≥4) who we defined as having DBD+CU.^43^ We then identified 258 typically developing (TD) children with no clinical diagnoses or CU traits (i.e., scores=0) who were matched to our DBD+CU group on age, sex, and income using a propensity score matching approach (see **eMethods 1** for more information). Consistent with prior ABCD studies, we excluded participants based on image quality for T1 and T2 images.^44^ These steps resulted in a final sample of 222 DBD+CU and 234 TD participants (**Table 1**).

**Table 1.** Demographic characteristics of the sample by subgroup.

|  | **Total (n=456)** | **TD (n=234)** | **S1 (n=84)** | **S2 (n=138)** | **P-value** |
| --- | --- | --- | --- | --- | --- |
| **Age** | 9.95 (0.62) | 9.94 (0.63) | 9.95 (0.66) | 9.98 (0.59) | 0.9 |
| **Sex** |  |  |  |  | >0.9 |
| Male | 324 (71%) | 167 (71%) | 59 (70%) | 98 (71%) |  |
| Female | 132 (29%) | 67 (29%) | 25 (30%) | 40 (29%) |  |
| **Race** |  |  |  |  | **0.025** |
| White | 285 (63%) | 150 (64%) | 59 (70%) | 76 (55%) |  |
| Black (African American) | 85 (19%) | 38 (16%) | 9 (11%) | 38 (28%) |  |
| Asian | 8 (1.8%) | 7 (3.0%) | 0 (0%) | 1 (0.7%) |  |
| American Indian/Alaska Native | 3 (0.7%) | 1 (0.4%) | 0 (0%) | 2 (1.4%) |  |
| Native Hawaiian or Other Pacific Islander | 2 (0.4%) | 2 (0.9%) | 0 (0%) | 0 (0%) |  |
| More than One Race | 55 (12%) | 24 (10%) | 13 (15%) | 18 (13%) |  |
| Unknown or Not Reported | 18 (3.9%) | 12 (5.1%) | 3 (3.6%) | 3 (2.2%) |  |
| **Household Income** |  |  |  |  | 0.12 |
| Less than \$11,999 | 49 (21%) | 24 (29%) | 41 (30%) | 114 (25%) | |
| \$12,000 - \$25,000 | 25 (11%) | 12 (14%) | 22 (16%) | 59 (13%) | |
| \$25,000 - \$49,000 | 48 (21%) | 13 (15%) | 34 (25%) | 95 (21%) | |
| \$50,000 - \$99,000 | 64 (27%) | 17 (20%) | 23 (17%) | 104 (23%) | |
| \$100,000 and greater | 24 (10%) | 8 (9.5%) | 8 (5.8%) | 40 (8.8%) | |
| Declined to answer or Don't Know | 24 (10%) | 10 (12%) | 10 (7.2%) | 44 (9.6%) |  |

Note: **TD** = typically developing; **S1** = subtype 1; **S2** = subtype 2

### Imaging Measures

#### Acquisition

Details of the imaging acquisition protocol have been described previously.^41^ Briefly, participants completed a 90-120 minute MRI scan including localizer, T1-weighted and T2-weighted scans. Imaging parameters differed slightly across sites, due to manufacturer differences (e.g., Siemens Prisma, Phillips, GE 750 3T scanners). All imaging data were processed by the ABCD Data Core^49^ and accessed as part of the 6.0 data release.

#### Structural MRI Preprocessing and Brain Segmentation

We used tabulated region of interest (ROI) data available from the National Institute for Mental Health Data Archive. Quality control procedures included both manual review as well as automated processes for T1– and T2– weighted images. sMRI underwent gradient nonlinearity, bias field, and intensity inhomogeneity corrections before being registered and resampled into an averaged, study specific reference brain in standard space. Cortical and subcortical segmentation were performed using FreeSurfer v5.3, using the Desikan-Killiany Atlas (cGMV) and the automatic subcortical segmentation of a brain volume (sGMV).

### Clinical Phenotyping and Environmental Measures

We evaluated the construct validity of groups using additional, independent measures. First, we evaluated externalizing psychopathology across groups using parent-reported aggression on the CBCL aggression subscale at baseline and parent reports of CU traits on 18 of the 24 original items from the Inventory of Callous Unemotional Traits (ICU),^45^ which was collected at 5 of the 21 ABCD sites at 2-year follow-up as part of the ABCD Social Development Substudy (ABCD-SD)^46^ (**eMethods 2**). Second, we evaluated how groups differed on other clinical and temperament constructs hypothesized to differentiate primary and secondary CU traits, including symptoms of anxiety and depression, which were assessed using child-report on the K-SADS at baseline^42^ and levels of fear and affiliation, which were assessed using parent report on the Early Adolescent Temperament Questionnaire (EATQ)^47^ at the 2-year follow-up. Third, we compared groups on exposomic risk using a previously validated measure of environmental risk and protective factors generated from a series of iterative bifactor analyses characterizing exposure to 348 factors across family, neighborhood, school, and state levels, including prenatal exposure to drugs, neighborhood poverty, and experience of discrimination.^4844^ Higher scores indicated greater exposure to adversity.

### Analysis

#### Data Harmonization

To account for site and scanner effects in imaging data, we implemented Longitudinal ComBat,^50^ leveraging a generalized additive mixed effect modeling strategy to allow for nonlinear effects of age (**eFigure 1**). ROIs for both cortical and subcortical GM were harmonized, preserving effects of age, sex, diagnostic status, and intracranial volume.

**Figure 1.**
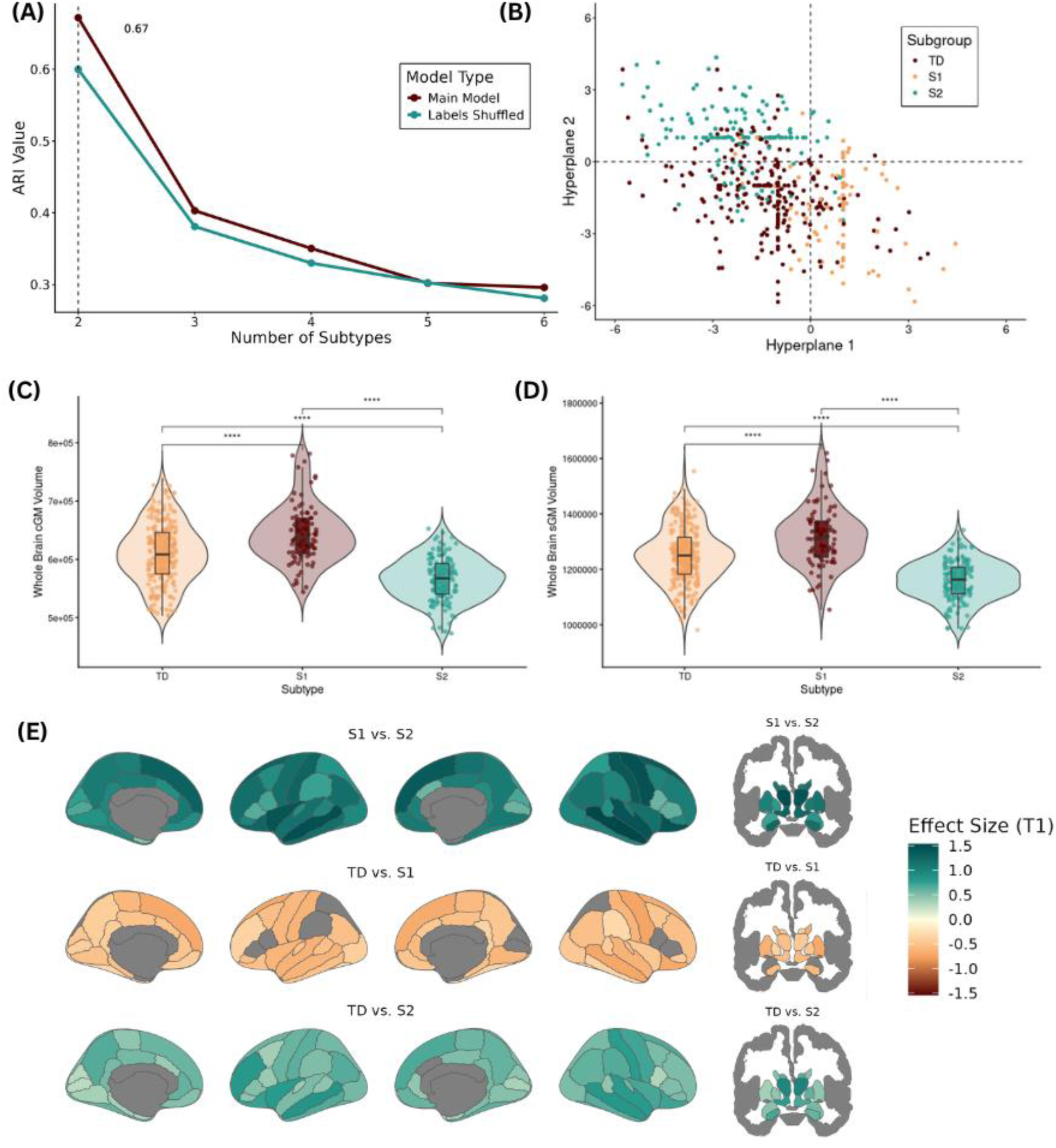
Heterogeneity through Discriminative Analysis (HYDRA) subtype identification. **(A)** The stability of the clustering solution after cross-validation was evaluated over a resolution range of 2–6 clusters and was quantified by the adjusted rand index (ARI). The maximum ARI (0.67) was seen with 2 subtypes, which also evidenced the most significant permutation test when compared to models run with shuffled labels. **(B)** The HYDRA model tried to achieve maximum separation between TD (n=234) and DBD/CU brains (n=222), while making 2 dimensions of CU traits (Hyperplane 1 [Primary] and Hyperplane 2 [Secondary]) most distinct. **(C, D)** S1 evidenced globally higher cGM and sGM than TD and S2, while S2 evidenced globally lower cGM and sGM. **(E)** Typically developing, S1, and S2 subgroups each differed significantly from each other in regional cGM and sGM across ROIs.

#### Weakly-Supervised Clustering

We implemented Heterogeneity through Discriminative Analysis (HYDRA), a weakly-supervised machine learning algorithm, to identify neurostructural subtypes of DBD+CU.^51,52^ HYDRA leverages multiple hyperplanes to distinguish between potential clinical subtypes using their differences from control populations (see **eMethods 3, eFigure 2**). Analyses were implemented in Python using the Machine Learning in Neuroimaging package (https://github.com/anbai106/mlni). HYDRA defined subtypes using cortical and subcortical GMV from 84 regions of interest, adjusting for age, sex, income, and intracranial volume. Consistent with other studies,^38–40^ we derived multiple solutions, requesting 2-6 clusters. We evaluated the stability of each solution using the Adjusted Rand Index, which was calculated using 10-fold cross validation. Finally, we conducted permutation testing (100 runs, TD vs. CU labels shuffled) to assess how each clustering solution differed from chance. To evaluate the stability of subtype assignment over time, we applied the trained model (e.g., the hyperplanes generated from the HYDRA analysis) to regional GMV values collected at 2-year follow-up (see **eMethods 4**).

**Figure 2.**
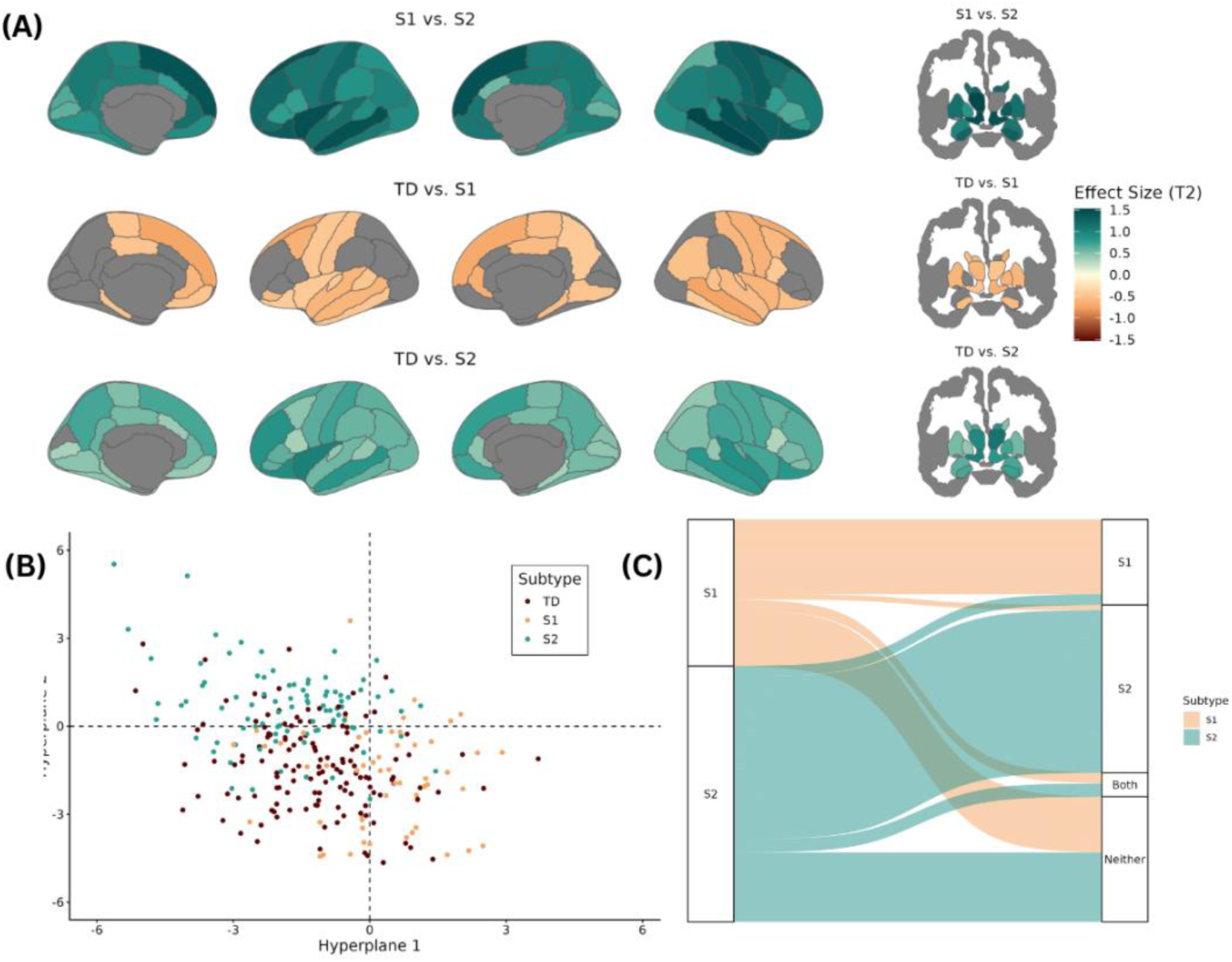
Subtypes identified by HYDRA remained stable over time. (**A**) Standardized effect sizes from ANOVA models at T3 show subtypes continue to differ from one another in their GMV across several cortical and subcortical ROIs. (**B**) The HYDRA model applied to 2-year follow-up data once again tried to achieve maximum separation between TD, S1, and S2 brains. (**C**) New subtype labels were assigned based on whether the SVM decision scores were positive (e.g., assumed to be associated with that subtype). Two S1 participants changed classification from S1 to S2, and 25 participants were assigned to ‘both’ or ‘neither’. Four S2 participants changed classification from S2 to S1, while 31 were assigned to both or neither subtype.

#### Group Level Statistical Analysis

All group level analyses were performed in R. We used analysis of covariance (ANCOVA) to evaluate the differences between groups on relevant clinical constructs, temperament and environmental exposures, thus providing construct validity tests of the identified neuroanatomical subtypes. Significant omnibus effects were followed by pairwise *post hoc* comparisons using Tukey-adjusted estimated marginal means. Multiple comparisons were corrected for across model families using the False Discovery Rate. All models retained age and sex as covariates.

## Results

### Two neuroanatomical subtypes of CU traits

HYDRA models identified between 2 and *k* neuroanatomical subtypes from 84 regional brain features (cGM and sGM), controlling for age, sex, household income, and ICV. Evaluation of cluster stability exhibited a clear peak at *k*=2 clusters with the most significant permutation testing (**Figure 1A**), suggesting the existence of 2 highly reproducible subtypes within DBD+CU (ARI=.67). Subtype 1 (*N*=84), delineated by Hyperplane 1, will hereafter be described as “S1” while Subtype 2 (*N*=138), delineated by Hyperplane 2 will hereafter be described as “S2” (**Figure 1B**).

Children in S1 were characterized by a widespread pattern of higher cGMV and sGMV than both S2 and TD children. *Post hoc* comparisons revealed that S1 evidenced greater GM than TD in 68 of 84 regions, and greater GMV than S2 in 83 of 84 regions (**eTable 1, Figure 1C**).

Conversely, children in S2 exhibited lower GMV than TD and S1 children. *Post hoc* comparisons revealed that S2 evidenced lower GMV than the TD group in 83 out of 84 regions (**eTable 1, Figure 1C**). Re-running analyses using the cGMV and sGMV data collected at 2 year follow-up demonstrated that subtypes continued to evidence widespread differences in regional GMV (**eTable 2, Figure 2A**). Subtype assignment remained relatively stable at 2-year follow-up **(Figure 2B)**, with subtype assignment at baseline significantly associated with subtype assignment at 2-year follow-up, χ²=19.185, *p*<.001 (**Figure 2C**). However, S2 were more likely to receive the same label at 2-year follow-up than S1 (χ²=67.76, *p*<.001, **eFigure 5**).

### Validation Using Clinical and Environmental Risk Indicators

Both S1 and S2 evidenced higher CU traits than TD children (*F*(2,451)=50.62, *p*<.001) but did not differ from each other (**Figure S4**). Likewise, both S1 and S2 were reported by parents to show higher aggression than TD children at baseline (*F*(2,451)=415.68, *p*<.001) but did not differ significantly from each other (**Figure 3A)**. However, subtypes did differ significantly based on self-reported anxiety symptoms (*F*(2,451)=7.43; *p*< .001; **Table 2**).

**Figure 3.**
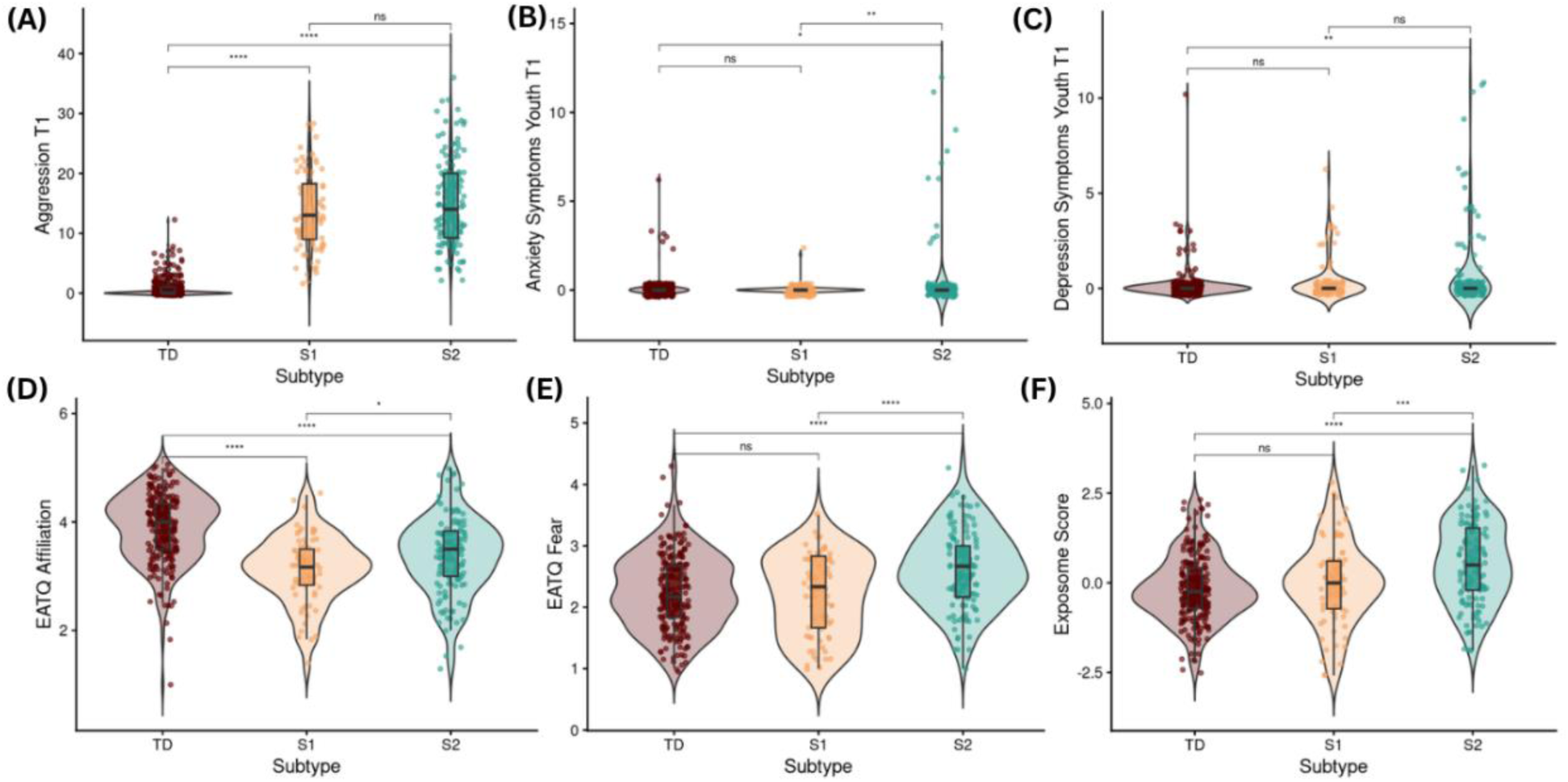
Neuroanatomical subtypes differ on relevant clinical, environmental, and temperamental factors. **(A)** Groups differed significantly on measures of parent report of aggression (F(2,451) = 48.0334, p<.001), and post-hoc tests revealed that both CU subtypes evidencing higher levels of aggression than the TD group but did not differ significantly from each other. **(B)** Groups differed significantly on child report of anxiety symptoms (F(2,451) = 7.43, p<.001) and post hoc comparisons revealed S2 had higher levels of anxiety than S1. (**C)** While groups differed significantly on child report of depression symptoms, post-hoc comparisons revealed that S1 and S2 did not significantly differ from each other. **(D)** Groups differed significantly on measures of affiliative temperament (F(2,451) = 48.03, p<.001), and post-hoc tests revealed that both CU subtypes evidenced lower levels of affiliation than the TD group. **(E)** Groups differed significantly on measures of fearful temperament (F(2,451) = 17.30, p<.001), and post-hoc tests revealed that S2 evidenced higher levels of fearfulness than S1. **(F)** Groups differed significantly on adversity exposure (F(2,451) = 19.34, p<.001) and post hoc tests revealed that S2 had higher exposome scores than S1.

**Table 2.**
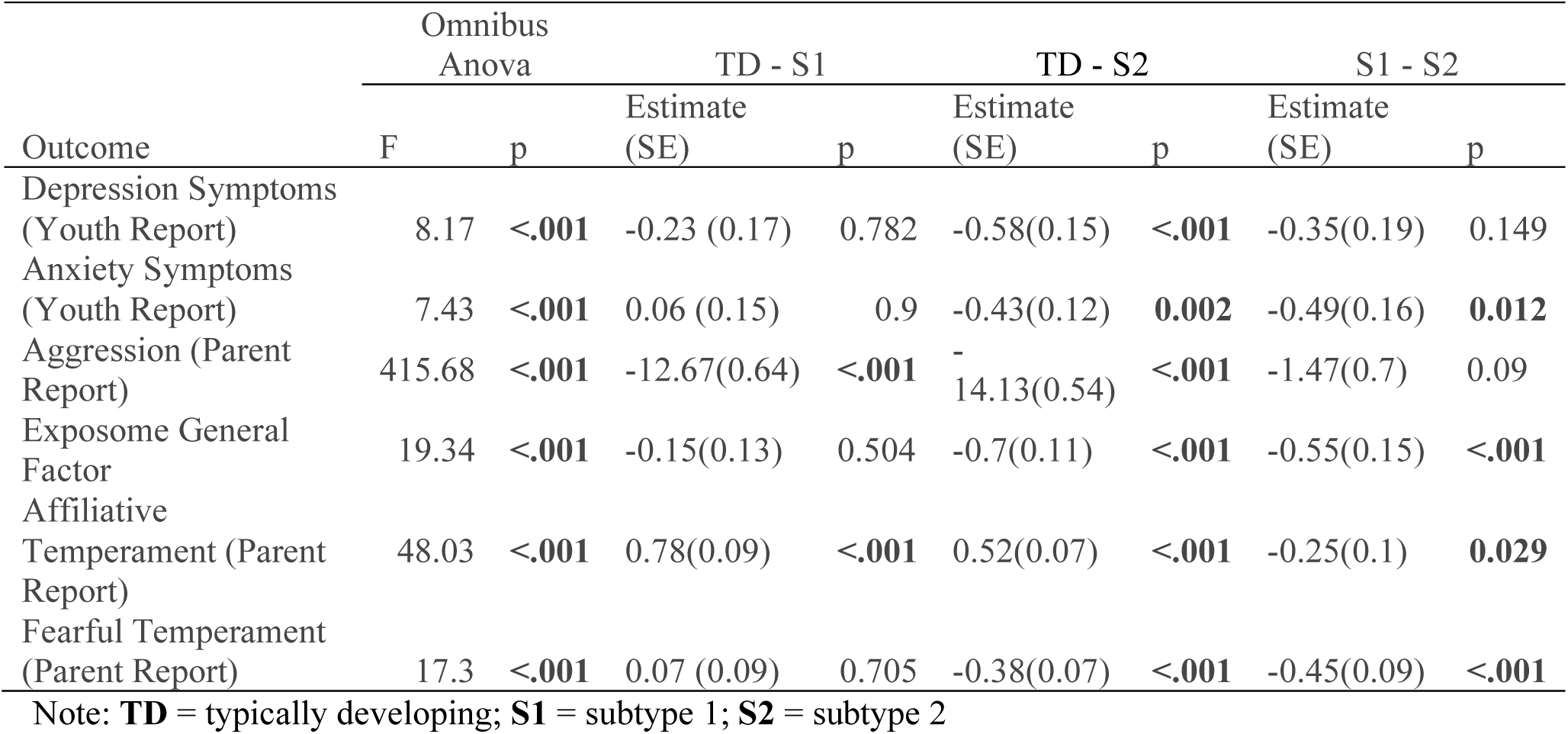
Comparisons across subtypes of clinical, environmental, and temperamental outcomes.

| Outcome | Omnibus<br>Anova |  | TD - S1 |  | TD - S2 |  | S1 - S2 |  |
| --- | --- | --- | --- | --- | --- | --- | --- | --- |
|  | F | p | Estimate<br>(SE) | p | Estimate<br>(SE) | p | Estimate<br>(SE) | p |
| Depression Symptoms<br>(Youth Report) | 8.17 | <.001 | -0.23 (0.17) | 0.782 | -0.58(0.15) | <.001 | -0.35(0.19) | 0.149 |
| Anxiety Symptoms<br>(Youth Report) | 7.43 | <.001 | 0.06 (0.15) | 0.9 | -0.43(0.12) | **0.002** | -0.49(0.16) | **0.012** |
| Aggression (Parent<br>Report) | 415.68 | <.001 | -12.67(0.64) | <.001 | -14.13(0.54) | <.001 | -1.47(0.7) | 0.09 |
| Exposome General<br>Factor | 19.34 | <.001 | -0.15(0.13) | 0.504 | -0.7(0.11) | <.001 | -0.55(0.15) | <.001 |
| Affiliative<br>Temperament (Parent<br>Report) | 48.03 | <.001 | 0.78(0.09) | <.001 | 0.52(0.07) | <.001 | -0.25(0.1) | **0.029** |
| Fearful Temperament<br>(Parent Report) | 17.3 | <.001 | 0.07 (0.09) | 0.705 | -0.38(0.07) | <.001 | -0.45(0.09) | <.001 |

Note: **TD** = typically developing; **S1** = subtype 1; **S2** = subtype 2

Specifically, S2 exhibited higher levels of anxiety than both S1 (*p*<.05) and TD children (*p*<.01), though S1 and TD did not differ from each other (**Figure 3B**). Subtypes also differed significantly on symptoms of depression (F(2,451) = 8.17: *p*<.001, **Table 2**), with S2 exhibiting higher levels of depression than TD children (*p* <.001). There was no difference between S1 and TD or S2 (**Figure 3C**). Affiliative temperament also varied by subtype (*F*(2,451) = 48.03, *p*<.001), with both S1 and S2 exhibiting lower affiliation than TD children (both *p*s < .001), and S1 exhibiting lower affiliation than S2 (*p*<.05, **Figure 3D**). Fearful temperament likewise differed between subgroups (*F*(2,451)=17.30, *p*<.001), with S2 showing greater fearfulness than both TD (*p*<.001) and S1 (*p*<.001), whereas TD and S1 did not differ (**Figure 3E**). Finally, exposome scores differed across groups (*F*(2,451)=19.34, *p*<.001), with S2 exhibiting higher exposome scores than both TD and S1 (both *ps*<.001), whereas TD and S1 did not differ (**Figure 3F**).

## Discussion

We used weakly-supervised machine learning methods to identify two reliable neuroanatomical subtypes of children with DBD+CU. The first subtype, S1, was characterized by higher cGM and sGM than both S2 and TD youth, with especially pronounced differences in superior frontal regions. Clinically, S1 showed fewer anxiety symptoms and had experienced less environmental adversity. In contrast, S2 was characterized by lower cGM and sGM than both S1 and TD youth, alongside more anxiety symptoms, greater environmental adversity, and a more fearful temperament. The identified DBD+CU subtypes were relatively stable over time, with similar patterns of cGM and sGM evidenced at the 2-year follow-up. Together, these findings suggest that clinically meaningful heterogeneity among children with CU traits is reflected in distinct neuroanatomical profiles.

These profiles may help explain inconsistent findings in prior neuroimaging research on CU traits and DBD. For example, reduced GMV has historically been associated with risk for psychopathology,^53–55^ although findings for externalizing psychopathology have been mixed.^56–58^ A recent meta-analysis found that DBDs were associated with greater *variability* in GM volumes, suggesting substantial heterogeneity in the neurobiological substrates of antisocial behavior.^59^ Our findings support this interpretation by showing that children with DBD+CU exhibit divergent patterns of GMV, with one subgroup showing relatively higher GMV and another showing relatively lower GMV. Likewise, reports that uncaring traits were associated with increased GMV in the orbitofrontal and anterior cingulate cortices^16^ may be explained by a greater proportion of children resembling our S1 profile. Conversely, other studies that reported reduced GMV in similar regions may be driven samples comprising children resembling our S2 profile,^60^ whose neuroanatomical pattern is more consistent with profiles observed among children who have experienced greater adversity, including deprivation and threat.^61^ Similar subtype differences could also explain inconsistent findings linking amygdala structure to both CU traits and adult psychopathy.^62^

Importantly, subtypes continued to differ from each other in GMV across most cortical and subcortical ROIs at 2-year follow up. However, differences between TD children and S1 were less pronounced, particularly in prefrontal regions. One possibility is that S1 reflects delayed cortical maturation rather than a stable pattern of elevated GMV.^58,63,64^ This explanation is consistent with our finding that subtype assignment was less stable over time for S1 than S2. In contrast, the lower GMV profile observed in S2 could reflect accelerated cortical maturation associated with experiences of early life stress or adversity.^65,66^ However, future research is needed to examine longitudinal trajectories of brain development across a wider age range.

The clinical correlates of S1 and S2 identified in the current work align with a longstanding theories of primary and secondary CU traits and psychopathy.^18,6718,68–71^ Both subtypes exhibited equally high levels of aggression and CU traits on independent measures, supporting convergent validity. However, our S1 subtype most closely mirrored conceptualizations of primary CU traits, being characterized by fewer internalizing symptoms than S2, lower fear and affiliation than both S2 and TD, and less exposure to environmental adversity than S2 subgroup. In contrast, S2 was characterized by more symptoms of anxiety, higher levels of fear, and greater exposomic risk. Together, these findings suggest that primary and secondary CU traits may reflect partially distinct etiological pathways, and that models that treat CU traits as unidimensional construct may obscure important biological and clinical differences, as well as treatment needs.

### Limitations

Several limitations of the current study should be considered. First, although our measure of CU traits exhibits strong psychometric properties^43^ and has been widely used in both neuroimaging and behavioral research,^72,73^ it comprises only four items, including three reverse-coded items from a prosocial behavior measure. At the same time, S1 and S2 differed from TD youth, but not each other, on a fuller, widely validated measures of CU traits at 2-year follow-up This pattern provides support for the convergent validity of our DBD+CU classification while also suggesting that the neuroanatomical subtypes we identified are unlikely to reflect differences in severity of CU traits. Second, the ABCD study recruited a community sample with a lower incidence of severe externalizing psychopathology, motivating our analytic decision to identify a smaller subsample that met diagnostic criteria for CD and ODD. Nevertheless, future studies are needed in samples recruited from clinical or juvenile justice settings to maximize generalizability. Finally, given the design of the ABCD study, we included some analyses using measures that had only been collected at follow-up timepoints. For example, our measure of temperament was collected at the 2-year follow up timepoint, which introduces a temporal mismatch between the psychological and neuroanatomical data used to generate and validate the subtypes. Additionally, some measures included in the exposome score were collected at the 1-year follow up assessment.^48^ However, many exposome features were retrospective in nature (e.g., birth complications) or drawn from geocoded items derived from baseline addresses.^74^ Further, prior work has demonstrated that measures of culture and environment in ABCD between baseline and 1-year follow-up demonstrate moderate to good test-retest reliability, suggesting that these variables may be relatively stable across this time.^75^

In conclusion, we identified two neuroanatomical subtypes of children with DBD+CU that differed in GMV, anxiety, adversity exposure, and temperament. These findings suggest that clinical distinctions between primary and secondary CU traits may have a detectable neurobiological basis. More broadly, our results highlight the importance of precision approaches to CU traits that move beyond average group differences and instead identify biologically and clinically meaningful heterogeneity within the phenotype. Such approaches may improve early identification, clarify developmental mechanisms, and inform more targeted interventions for children with DBD and CU traits, who are at risk for severe lifespan trajectories of antisocial conduct.^19,76^

## Acknowledgments

Data used in the preparation of this article were obtained from the Adolescent Brain Cognitive Development (ABCD) Study (https://abcdstudy.org), held in the NIMH Data Archive (NDA). The ABCD Study is supported by the National Institutes of Health and additional federal partners. This manuscript reflects the views of the authors and may not reflect the opinions or views of the NIH or ABCD consortium investigators. The preparation of this manuscript was also partially supported by funding from the National Institute of Mental Health to Murtha (1F31MH140550-01) Waller (R01MH125904) and Seidlitz (R01MH133843).

## References

1. Frick PJ, Ellis M. Callous-Unemotional Traits and Subtypes of Conduct Disorder. Clin Child Fam Psychol Rev. 1999;2(3):149–168. doi:10.1023/A:1021803005547

2. Waller R, Wagner NJ, Barstead MG, et al. A meta-analysis of the associations between callous-unemotional traits and empathy, prosociality, and guilt. Clinical Psychology Review. 2020;75:101809. doi:10.1016/j.cpr.2019.101809

3. Frick PJ, White SF. Research Review: The importance of callous-unemotional traits for developmental models of aggressive and antisocial behavior. Journal of Child Psychology and Psychiatry. 2008;49(4):359–375. doi:10.1111/j.1469-7610.2007.01862.x

4. Fairchild G, Hawes DJ, Frick PJ, et al. Conduct disorder. Nat Rev Dis Primers. 2019;5(1):43. doi:10.1038/s41572-019-0095-y

5. Goulter N, Hur YS, Jones DE, et al. Kindergarten conduct problems are associated with monetized outcomes in adolescence and adulthood. Journal of Child Psychology and Psychiatry. n/a(n/a). doi:10.1111/jcpp.13837

6. Rivenbark JG, Odgers CL, Caspi A, et al. The high societal costs of childhood conduct problems: evidence from administrative records up to age 38 in a longitudinal birth cohort. Journal of Child Psychology and Psychiatry. 2018;59(6):703–710. doi:10.1111/jcpp.12850

7. Todorov JJ, Devine RT, De Brito SA. Association between childhood maltreatment and callous-unemotional traits in youth: A meta-analysis. Neuroscience & Biobehavioral Reviews. 2023;146:105049. doi:10.1016/j.neubiorev.2023.105049

8. Moore AA, Blair RJ, Hettema JM, Roberson-Nay R. The genetic underpinnings of callous-unemotional traits: A systematic research review. Neuroscience & Biobehavioral Reviews. 2019;100:85–97. doi:10.1016/j.neubiorev.2019.02.018

9. Blair RJR, Zhang R. Recent neuro-imaging findings with respect to conduct disorder, callous-unemotional traits and psychopathy. Curr Opin Psychiatry. 2020;33(1):45–50. doi:10.1097/YCO.0000000000000559

10. Wagner NJ, Waller R. Leveraging parasympathetic nervous system activity to study risk for psychopathology: The special case of callous-unemotional traits. Neuroscience & Biobehavioral Reviews. 2020;118:175–185. doi:10.1016/j.neubiorev.2020.07.029

11. Cardinale EM, O’Connell K, Robertson EL, et al. Callous and uncaring traits are associated with reductions in amygdala volume among youths with varying levels of conduct problems. Psychol Med. 2019;49(9):1449–1458. doi:10.1017/S0033291718001927

12. Dotterer HL, Waller R, Hein TC, et al. Clarifying the Link Between Amygdala Functioning During Emotion Processing and Antisocial Behaviors Versus Callous-Unemotional Traits Within a Population-Based Community Sample. Clin Psychol Sci. 2020;8(5):918–935. doi:10.1177/2167702620922829

13. Viding E, Sebastian CL, Dadds MR, et al. Amygdala Response to Preattentive Masked Fear in Children With Conduct Problems: The Role of Callous-Unemotional Traits. AJP. 2012;169(10):1109–1116. doi:10.1176/appi.ajp.2012.12020191

14. Paz Y, Perkins ER, Colins O, et al. Evaluating the sensitivity to threat and affiliative reward (STAR) model in relation to the development of conduct problems and callous-unemotional traits across early adolescence. J Child Psychol Psychiatry. Published online March 13, 2024. doi:10.1111/jcpp.13976

15. Waller R, Wagner N. The Sensitivity to Threat and Affiliative Reward (STAR) model and the development of callous-unemotional traits. Neurosci Biobehav Rev. 2019;107:656–671. doi:10.1016/j.neubiorev.2019.10.005

16. Caldwell BM, Anderson NE, Harenski KA, et al. The structural brain correlates of callous-unemotional traits in incarcerated male adolescents. Neuroimage Clin. 2019;22:101703. doi:10.1016/j.nicl.2019.101703

17. De Brito SA, Mechelli A, Wilke M, et al. Size matters: Increased grey matter in boys with conduct problems and callous–unemotional traits. Brain. 2009;132(4):843–852. doi:10.1093/brain/awp011

18. Craig SG, Goulter N, Moretti MM. A Systematic Review of Primary and Secondary Callous-Unemotional Traits and Psychopathy Variants in Youth. Clin Child Fam Psychol Rev. 2021;24(1):65–91. doi:10.1007/s10567-020-00329-x

19. Fleming GE, Neo B, Kaouar S, Kimonis ER. Treatment Outcomes of Children with Primary Versus Secondary Callous-Unemotional Traits. Res Child Adolesc Psychopathol. 2023;51(11):1581–1594. doi:10.1007/s10802-023-01112-6

20. Karpman B. On the need of separating psychopathy into two distinct clinical types: the symptomatic and the idiopathic. Journal of Criminal Psychopathology. 1941;3:112–137.

21. Cleckley H. The Mask of Sanity; an Attempt to Reinterpret the so-Called Psychopathic Personality. Mosby; 1941:298.

22. Kimonis ER. The Emotionally Sensitive Child-Adverse Parenting Experiences-Allostatic (Over)Load (ESCAPE-AL) Model for the Development of Secondary Psychopathic Traits. Clin Child Fam Psychol Rev. 2023;26(4):1097–1114. doi:10.1007/s10567-023-00455-2

23. Fanti KA, Kyranides MN, Petridou M, Demetriou CA, Georgiou G. Neurophysiological markers associated with heterogeneity in conduct problems, callous unemotional traits, and anxiety: Comparing children to young adults. Developmental Psychology. 2018;54(9):1634–1649. doi:10.1037/dev0000505

24. Kimonis ER, Skeem JL, Cauffman E, Dmitrieva J. Are secondary variants of juvenile psychopathy more reactively violent and less psychosocially mature than primary variants? Law Hum Behav. 2011;35(5):381–391. doi:10.1007/s10979-010-9243-3

25. Kimonis ER, Goulter N, Hawes DJ, Wilbur RR, Groer MW. Neuroendocrine factors distinguish juvenile psychopathy variants. Developmental Psychobiology. 2017;59(2):161–173. doi:10.1002/dev.21473

26. Kimonis ER, Fanti KA, Goulter N, Hall J. Affective startle potentiation differentiates primary and secondary variants of juvenile psychopathy. Development and Psychopathology. 2017;29(4):1149–1160. doi:10.1017/S0954579416001206

27. Dugré JR, Potvin S. Functional Connectivity of the Nucleus Accumbens across Variants of Callous-Unemotional Traits: A Resting-State fMRI Study in Children and Adolescents. Res Child Adolesc Psychopathol. 2024;52(3):353–368. doi:10.1007/s10802-023-01143-z

28. Fanti KA, Konikou K, Cohn M, Popma A, Brazil IA. Amygdala functioning during threat acquisition and extinction differentiates antisocial subtypes. J Neuropsychol. 2020;14(2):226–241. doi:10.1111/jnp.12183

29. Sethi A, McCrory E, Puetz V, et al. ‘Primary’ and ‘secondary’ variants of psychopathy in a volunteer sample are associated with different neurocognitive mechanisms. Biol Psychiatry Cogn Neurosci Neuroimaging. 2018;3(12):1013–1021. doi:10.1016/j.bpsc.2018.04.002

30. De Brito SA, Viding E, Sebastian CL, et al. Reduced orbitofrontal and temporal grey matter in a community sample of maltreated children. Journal of Child Psychology and Psychiatry. 2013;54(1):105–112. doi:10.1111/j.1469-7610.2012.02597.x

31. Liu X, Klugah-Brown B, Zhang R, Chen H, Zhang J, Becker B. Pathological fear, anxiety and negative affect exhibit distinct neurostructural signatures: evidence from psychiatric neuroimaging meta-analysis. Transl Psychiatry. 2022;12(1):405. doi:10.1038/s41398-022-02157-9

32. Morey RA, Haswell CC, Hooper SR, De Bellis MD. Amygdala, Hippocampus, and Ventral Medial Prefrontal Cortex Volumes Differ in Maltreated Youth with and without Chronic Posttraumatic Stress Disorder. Neuropsychopharmacol. 2016;41(3):791–801. doi:10.1038/npp.2015.205

33. Kimonis ER, Frick PJ, Cauffman E, Goldweber A, Skeem J. Primary and secondary variants of juvenile psychopathy differ in emotional processing. Development and Psychopathology. 2012;24(3):1091–1103. doi:10.1017/S0954579412000557

34. Campbell DT, Fiske DW. Convergent and discriminant validation by the multitrait-multimethod matrix. Psychological Bulletin. 1959;56(2):81–105. doi:10.1037/h0046016

35. Joyner KJ, Perkins ER. Challenges and ways forward in bridging units of analysis in clinical psychological science. Journal of Psychopathology and Clinical Science. 2023;132(7):888–896. doi:10.1037/abn0000879

36. Cuthbert BN, Insel TR. Toward the future of psychiatric diagnosis: the seven pillars of RDoC. BMC Medicine. 2013;11(1):126. doi:10.1186/1741-7015-11-126

37. Kaczkurkin AN, Moore TM, Sotiras A, Xia CH, Shinohara RT, Satterthwaite TD. Approaches to Defining Common and Dissociable Neurobiological Deficits Associated With Psychopathology in Youth. Biological Psychiatry. 2020;88(1):51–62. doi:10.1016/j.biopsych.2019.12.015

38. Baller EB, Kaczkurkin AN, Sotiras A, et al. Neurocognitive and functional heterogeneity in depressed youth. Neuropsychopharmacol. 2021;46(4):783–790. doi:10.1038/s41386-020-00871-w

39. Hwang G, Wen J, Sotardi S, et al. Assessment of Neuroanatomical Endophenotypes of Autism Spectrum Disorder and Association With Characteristics of Individuals With Schizophrenia and the General Population. JAMA Psychiatry. 2023;80(5):498–507. doi:10.1001/jamapsychiatry.2023.0409

40. Kaczkurkin AN, Sotiras A, Baller EB, et al. Neurostructural Heterogeneity in Youths With Internalizing Symptoms. Biological Psychiatry. 2020;87(5):473–482. doi:10.1016/j.biopsych.2019.09.005

41. Casey BJ, Cannonier T, Conley MI, et al. The Adolescent Brain Cognitive Development (ABCD) study: Imaging acquisition across 21 sites. Developmental Cognitive Neuroscience. 2018;32:43–54. doi:10.1016/j.dcn.2018.03.001

42. Kaufman J, Birmaher B, Brent D, et al. Schedule for Affective Disorders and Schizophrenia for School-Age Children-Present and Lifetime Version (K-SADS-PL): initial reliability and validity data. J Am Acad Child Adolesc Psychiatry. 1997;36(7):980–988. doi:10.1097/00004583-199707000-00021

43. Hawes SW, Waller R, Thompson WK, et al. Assessing callous-unemotional traits: development of a brief, reliable measure in a large and diverse sample of preadolescent youth. Psychological Medicine. 2020;50(3):456–464. doi:10.1017/S0033291719000278

44. Keller AS, Moore TM, Luo A, et al. A general exposome factor explains individual differences in functional brain network topography and cognition in youth. Developmental Cognitive Neuroscience. 2024;66:101370. doi:10.1016/j.dcn.2024.101370

45. Frick PJ. Inventory of Callous–Unemotional Traits. Published online 2004. doi:10.1037/t62639-000

46. Hoffman EA, Clark DB, Orendain N, Hudziak J, Squeglia LM, Dowling GJ. Stress exposures, neurodevelopment and health measures in the ABCD study. Neurobiol Stress. 2019;10:100157. doi:10.1016/j.ynstr.2019.100157

47. Ellis LK, Rothbart M. Early Adolescent Temperament Questionnaire--Revised. Published online 2001. doi:10.1037/t07624-000

48. Moore TM, Visoki E, Argabright ST, et al. Modeling environment through a general exposome factor in two independent adolescent cohorts. Exposome. 2022;2(1):osac010. doi:10.1093/exposome/osac010

49. Hagler DJ, Hatton SeanN, Cornejo MD, et al. Image processing and analysis methods for the Adolescent Brain Cognitive Development Study. NeuroImage. 2019;202:116091. doi:10.1016/j.neuroimage.2019.116091

50. Beer JC, Tustison NJ, Cook PA, et al. Longitudinal ComBat: A method for harmonizing longitudinal multi-scanner imaging data. NeuroImage. 2020;220:117129. doi:10.1016/j.neuroimage.2020.117129

51. Varol E, Sotiras A, Davatzikos C. HYDRA: Revealing heterogeneity of imaging and genetic patterns through a multiple max-margin discriminative analysis framework. NeuroImage. 2017;145:346–364. doi:10.1016/j.neuroimage.2016.02.041

52. Wen J, Varol E, Sotiras A, et al. Multi-scale semi-supervised clustering of brain images: Deriving disease subtypes. Medical Image Analysis. 2022;75:102304. doi:10.1016/j.media.2021.102304

53. Li T, Wang L, Camilleri JA, et al. Mapping common grey matter volume deviation across child and adolescent psychiatric disorders. Neuroscience & Biobehavioral Reviews. 2020;115:273–284. doi:10.1016/j.neubiorev.2020.05.015

54. Durham EL, Jeong HJ, Moore TM, et al. Association of gray matter volumes with general and specific dimensions of psychopathology in children. Neuropsychopharmacology. 2021;46(7):1333–1339. doi:10.1038/s41386-020-00952-w

55. Kaczkurkin AN, Park SS, Sotiras A, et al. Evidence for Dissociable Linkage of Dimensions of Psychopathology to Brain Structure in Youths. AJP. 2019;176(12):1000–1009. doi:10.1176/appi.ajp.2019.18070835

56. Valera EM, Faraone SV, Murray KE, Seidman LJ. Meta-Analysis of Structural Imaging Findings in Attention-Deficit/Hyperactivity Disorder. Biological Psychiatry. 2007;61(12):1361–1369. doi:10.1016/j.biopsych.2006.06.011

57. Nakao T, Radua J, Rubia K, Mataix-Cols D. Gray Matter Volume Abnormalities in ADHD: Voxel-Based Meta-Analysis Exploring the Effects of Age and Stimulant Medication. AJP. 2011;168(11):1154–1163. doi:10.1176/appi.ajp.2011.11020281

58. Phillips NL, Sharpe BM, Hyatt CS, et al. Structural brain correlates of externalizing traits and symptoms in the IMAGEN sample. *Personality Disorders: Theory*, Research, and Treatment. 2025;16(1):43–56. doi:10.1037/per0000701

59. Tully J, Cross B, Gerrie B, et al. A systematic review and meta-analysis of brain volume abnormalities in disruptive behaviour disorders, antisocial personality disorder and psychopathy. Nat Mental Health. 2023;1(3):163–173. doi:10.1038/s44220-023-00032-0

60. Sebastian CL, De Brito SA, McCrory EJ, et al. Grey Matter Volumes in Children with Conduct Problems and Varying Levels of Callous-Unemotional Traits. J Abnorm Child Psychol. 2016;44:639–649. doi:10.1007/s10802-015-0073-0

61. McLaughlin KA, Weissman D, Bitrán D. Childhood Adversity and Neural Development: A Systematic Review. Annual Review of Developmental Psychology. 2019;1(Volume 1, 2019):277–312. 10.1146/annurev-devpsych-121318-084950

62. Deming P, Heilicher M, Koenigs M. How reliable are amygdala findings in psychopathy? A systematic review of MRI studies. Neuroscience & Biobehavioral Reviews. 2022;142:104875. doi:10.1016/j.neubiorev.2022.104875

63. Shi R, Xiang S, Jia T, et al. Investigating grey matter volumetric trajectories through the lifespan at the individual level. Nature Communications. 2024;15(1):5954. doi:10.1038/s41467-024-50305-0

64. Gogtay N, Thompson PM. Mapping gray matter development: Implications for typical development and vulnerability to psychopathology. Brain and Cognition. 2010;72(1):6–15. doi:10.1016/j.bandc.2009.08.009

65. Lupien SJ, McEwen BS, Gunnar MR, Heim C. Effects of stress throughout the lifespan on the brain, behaviour and cognition. Nature Reviews Neuroscience. 2009;10(6):434–445. doi:10.1038/nrn2639

66. Nweze T, Banaschewski T, Ajaelu C, et al. Trajectories of cortical structures associated with stress across adolescence: a bivariate latent change score approach. Journal of Child Psychology and Psychiatry. 2023;64(8):1159–1175. doi:10.1111/jcpp.13793

67. Almas I, Lordos A. A narrative review of psychopathy research: current advances and the argument for a qualitative approach. The Journal of Forensic Psychiatry & Psychology. 2025;36(3):356–406. doi:10.1080/14789949.2025.2456208

68. Kimonis ER, Frick PJ, Munoz LC, Aucoin KJ. Callous-unemotional traits and the emotional processing of distress cues in detained boys: Testing the moderating role of aggression, exposure to community violence, and histories of abuse. Development and Psychopathology. 2008;20(2):569–589. doi:10.1017/S095457940800028X

69. Craig SG, Moretti MM. Profiles of primary and secondary callous-unemotional features in youth: The role of emotion regulation. Development and Psychopathology. 2019;31(4):1489–1500. doi:10.1017/S0954579418001062

70. Blair RJR. The amygdala and ventromedial prefrontal cortex in morality and psychopathy. Trends in Cognitive Sciences. 2007;11(9):387–392. doi:10.1016/j.tics.2007.07.003

71. Blair RJR. Psychopathy, frustration, and reactive aggression: The role of ventromedial prefrontal cortex. British Journal of Psychology. 2010;101(3):383–399. doi:10.1348/000712609X418480

72. Umbach RH, Tottenham N. Callous-unemotional traits and reduced default mode network connectivity within a community sample of children. Development and Psychopathology. 2021;33(4):1170–1183. doi:10.1017/S0954579420000401

73. Murtha K, Perlstein S, Paz Y, et al. Callous-unemotional traits, cognitive functioning, and externalizing problems in a propensity-matched sample from the ABCD study. Journal of Child Psychology and Psychiatry. 2025;66(3):333–349. doi:10.1111/jcpp.14062

74. Fan CC, Marshall A, Smolker H, et al. Adolescent Brain Cognitive Development (ABCD) study Linked External Data (LED): Protocol and practices for geocoding and assignment of environmental data. Developmental Cognitive Neuroscience. 2021;52:101030. doi:10.1016/j.dcn.2021.101030

75. Gonzalez R, Thompson EL, Sanchez M, et al. An update on the assessment of culture and environment in the ABCD Study®: Emerging literature and protocol updates over three measurement waves. Developmental Cognitive Neuroscience. 2021;52. doi:10.1016/j.dcn.2021.101021

76. Perlstein S, Fair M, Hong E, Waller R. Treatment of childhood disruptive behavior disorders and callous-unemotional traits: a systematic review and two multilevel meta-analyses. J Child Psychol Psychiatry. Published online March 1, 2023. doi:10.1111/jcpp.13774

