## Supplemental Files for "Neuroanatomical Subtypes of Callous-Unemotional Traits in a Community Sample of Youth"

**eMethods 1.** Matching Procedure

Using the R package MatchIt, CU/DBD youth were matched on age, sex and income with typically developing youth. Participants were matched across groups using one-to-one nearest neighbor matching with Caliper distance = .10, with propensity scores estimated using logistic regression. Matching procedures yielded adequate balance, where standardized mean differences of covariates between groups were all <.10. This resulted in 258 CU/DBD and 258 typically developing youth.

**eMethods 2.** Measure of CU Traits

To validate our groups created using our 4-item measure of CU traits, we compared mean scores for parent report of 18 of the 24 original items from the Inventory of Callous–Unemotional Traits (ICU; Frick, 2004; e.g., "I do not feel remorseful when I do something wrong," "I feel bad or guilty when I do something wrong," "The feelings of others are unimportant to me"). Items were rated on a 4-point scale (0=not at all true, 3=definitely true). This measure was collected as part of the ABCD Social Development Substudy (ABCD-SD), for which data were collected at 5 of the 21 ABCD sites at 2-year follow-up (Hoffman et al., 2019). Of the subjects included in the clustering sample, 48 TD, 20 S1, and 28 S2 participants had data from the Social Development Substudy (**see Figure S3**). The 4 item measure of CU traits and measure from the ABCD-SD were strongly correlated, Pearson's R = 0.72, $t(1,94)=10.18$, $p<.001$.

**eMethods 3.** HYDRA

Heterogeneity through Discriminative Analysis (HYDRA) leverages support vector machines (SVM), which are widely used supervised machine learning methods used for discrimination tasks (Shmilovici, 2005). HYDRA applies multiple linear SVMs to non-linear classification cases to perform simultaneously clustering and classification. Specifically, it constructs a complex “polytope” (e.g., a geometric object with flat sides or face) combining the hyperplanes from *k* linear SVMs and separating typically developing participants from each of the *k* identified subgroups from the patient population (**Figure S1**). Each face of the convex polytope represents a unique subtype that reflects a specific clinical pattern within the input features from the patient population. A key strength of HYDRA and weakly-supervised clustering is their ability to minimize the influence of disease-unrelated confounds by focusing on clustering the differences between patients and controls.

**eMethods 4.** Applying HYRDA to external data

After the model was trained on the DBD/CU population, the optimal polytope which best discriminates the typically developing (TD) group from the *k* subtypes of DBD/CU is saved. The *k* sets of weights are then applied to the same 84 ROI's collected at 2-year follow-up, resulting in *k* SVM scores for each participant which represent the distance from each hyperplane (Wen et al., 2022). Therefore, larger scores represent greater separation from the TD group, where smaller scores represent less separation, or neuroanatomy that is more similar to that of a TD participant. If a participant receives a positive score for one hyperplane, they are predicted to belong to that subtype. If a participant receives a negative score for one hyperplane, they are predicted to not belong to that subtype. If a participant receives a positive score for both hyperplanes, they are assigned to the subtype with the larger score. If both scores are negative, they are assigned to the subtype with the smaller score.

**eTable 1.** Comparisons across subtypes of regional GM volumes

| | Omnibus Anova | | TD - S1 | | TD - S2 | | S1 - S2 | | Omnibus Anova | | TD - S1 | | TD - S2 | | S1 - S2 | |
|---|---|---|---|---|---|---|---|---|---|---|---|---|---|---|---|---|
| Region | F | p | Estimate (SE) | p | Estimate (SE) | p | Estimate (SE) | p | F | p | Estimate (SE) | p | Estimate (SE) | p | Estimate (SE) | p |
| | **Left Hemisphere** | | | | | | | | **Right Hemisphere** | | | | | | | |
| Nucleus Acumbens | 34.34 | **<.001** | -43.24(12.02) | **0.002** | 60.8(10.14) | **<.001** | 104.05(13.07) | **<.001** | 34.69 | **<.001** | -35.73(10.68) | **0.004** | 55.77(9.01) | **<.001** | 91.5(11.62) | **<.001** |
| Amygdala | 16.04 | **<.001** | -49.05(24.47) | 0.124 | 91.77(20.66) | **<.001** | 140.82(26.63) | **<.001** | 20.21 | **<.001** | -53.17(24.68) | 0.093 | 105.26(20.83) | **<.001** | 158.43(26.86) | **<.001** |
| Caudate | 32.25 | **<.001** | -267.12(55.14) | **<.001** | 213.43(46.54) | **<.001** | 480.55(60) | **<.001** | 35.17 | **<.001** | -285.23(53.59) | **<.001** | 204.04(45.23) | **<.001** | 489.27(58.32) | **<.001** |
| Hippocampus | 42.7 | **<.001** | -189.31(43.6) | **<.001** | 238.07(36.8) | **<.001** | 427.38(47.44) | **<.001** | 36.38 | **<.001** | -204.08(45.35) | **<.001** | 211.86(38.27) | **<.001** | 415.94(49.34) | **<.001** |
| Pallidum | 17.66 | **<.001** | -85.53(29.93) | **0.017** | 103.9(25.26) | **<.001** | 189.43(32.56) | **<.001** | 32.29 | **<.001** | -88.44(24.55) | **0.002** | 119.2(20.72) | **<.001** | 207.64(26.71) | **<.001** |
| Putamen | 26.15 | **<.001** | -337.42(66.7) | **<.001** | 191.99(56.3) | **0.002** | 529.41(72.58) | **<.001** | 33.92 | **<.001** | -354.85(64.25) | **<.001** | 224.69(54.23) | **<.001** | 579.54(69.91) | **<.001** |
| Thalamus | 65.43 | **<.001** | -334.32(80.66) | **<.001** | 606.27(68.08) | **<.001** | 940.59(87.77) | **<.001** | 64.92 | **<.001** | -244.57(71.15) | **0.003** | 558.11(60.05) | **<.001** | 802.68(77.42) | **<.001** |
| Ventral Diencephalon | 47.54 | **<.001** | -165.78(41.5) | **<.001** | 256.42(35.03) | **<.001** | 422.2(45.16) | **<.001** | 42.35 | **<.001** | -183.82(41.85) | **<.001** | 227.1(35.32) | **<.001** | 410.92(45.54) | **<.001** |
| Banks of the Superior Temporal Sulcus | 24.58 | **<.001** | -275.09(68.08) | **<.001** | 240.36(57.46) | **<.001** | 515.45(74.08) | **<.001** | 21.06 | **<.001** | -199.1(58.11) | **0.003** | 203.48(49.05) | **<.001** | 402.58(63.23) | **<.001** |
| Caudal Anterior Cingulate | 20.85 | **<.001** | -275.03(71.15) | **<.001** | 223.93(60.05) | **<.001** | 498.95(77.42) | **<.001** | 9.07 | **<.001** | -250.04(72.08) | **0.003** | 77.04(60.84) | 0.415 | 327.08(78.43) | **<.001** |
| Caudal Middle Frontal | 21.45 | **<.001** | -787.79(188.38) | **<.001** | 556.65(159) | **0.002** | 1344.44(204.99) | **<.001** | 27.16 | **<.001** | -668.78(181.78) | **0.001** | 748.37(153.43) | **<.001** | 1417.15(197.8) | **<.001** |
| Cuneus | 14.93 | **<.001** | -299.33(81.43) | **0.001** | 184.22(68.73) | **0.021** | 483.55(88.61) | **<.001** | 14.7 | **<.001** | -168.96(91.72) | 0.168 | 328.91(77.41) | **<.001** | 497.87(99.8) | **<.001** |
| Entorhinal | 6.65 | **0.001** | -45.37(49.5) | 0.645 | 128.33(41.78) | **0.007** | 173.7(53.86) | **0.004** | 4.8 | **0.009** | 25.14(48.52) | 0.862 | 125.88(40.95) | **0.007** | 100.73(52.8) | 0.138 |
| Fusiform | 26.25 | **<.001** | -439.25(172.57) | **0.038** | 821.19(145.65) | **<.001** | 1260.44(187.78) | **<.001** | 28.26 | **<.001** | -461.16(169.31) | **0.024** | 829.7(142.9) | **<.001** | 1290.86(184.23) | **<.001** |
| Isthmus Cingulate | 25.17 | **<.001** | -201.79(62.59) | **0.006** | 263.52(52.83) | **<.001** | 465.31(68.11) | **<.001** | 24 | **<.001** | -174.17(61.63) | **0.018** | 265.35(52.02) | **<.001** | 439.52(67.07) | **<.001** |
| Insula | 54.68 | **<.001** | -472.28(98.15) | **<.001** | 607.54(82.85) | **<.001** | 1079.81(106.81) | **<.001** | 38.59 | **<.001** | -347.29(96.81) | **0.002** | 532.16(81.71) | **<.001** | 879.45(105.34) | **<.001** |
| Inferior Parietal | 38.22 | **<.001** | -1086.87(269.04) | **<.001** | 1382.31(227.08) | **<.001** | 2469.17(292.75) | **<.001** | 35.12 | **<.001** | -1248.76(307.23) | **<.001** | 1474.04(259.31) | **<.001** | 2722.8(334.31) | **<.001** |
| Inferior Temporal | 38.02 | **<.001** | -898.08(222.65) | **<.001** | 1153.16(187.92) | **<.001** | 2051.24(242.27) | **<.001** | 29.79 | **<.001** | -613.44(224.66) | **0.024** | 1149.57(189.63) | **<.001** | 1763.01(244.47) | **<.001** |
| Lingual | 19.63 | **<.001** | -581.68(146.56) | **<.001** | 412.5(123.7) | **0.003** | 994.18(159.48) | **<.001** | 31.17 | **<.001** | -707.64(150.13) | **<.001** | 571.14(126.71) | **<.001** | 1278.78(163.36) | **<.001** |
| Lateral Orbitofrontal | 44.75 | **<.001** | -472.65(118.33) | **<.001** | 689.63(99.88) | **<.001** | 1162.28(128.76) | **<.001** | 54.95 | **<.001** | -629.13(113.61) | **<.001** | 645.4(95.89) | **<.001** | 1274.53(123.62) | **<.001** |
| Lateral Occipital | 25.65 | **<.001** | -701.1(225.17) | **0.008** | 975.36(190.05) | **<.001** | 1676.47(245.02) | **<.001** | 25.76 | **<.001** | -885.79(251.76) | **0.002** | 1032.02(212.5) | **<.001** | 1917.81(273.96) | **<.001** |
| Medial Orbitofrontal | 22.52 | **<.001** | -279.2(85.6) | **0.005** | 327.86(72.25) | **<.001** | 607.07(93.14) | **<.001** | 32.31 | **<.001** | -321.99(85.5) | **0.001** | 400.94(72.17) | **<.001** | 722.93(93.04) | **<.001** |
| Middle Temporal | 60.11 | **<.001** | -987.64(203.92) | **<.001** | 1352.56(172.12) | **<.001** | 2340.19(221.9) | **<.001** | 64.42 | **<.001** | -1165.47(206.8) | **<.001** | 1332.32(174.55) | **<.001** | 2497.79(225.03) | **<.001** |
| Paracentral | 27.64 | **<.001** | -327.38(71.77) | **<.001** | 250.93(60.57) | **<.001** | 578.31(78.09) | **<.001** | 35.11 | **<.001** | -361.12(80.54) | **<.001** | 360.69(67.98) | **<.001** | 721.8(87.65) | **<.001** |
| Pericalcarine | 12.04 | **<.001** | -191.94(61.25) | **0.007** | 134.04(51.7) | **0.027** | 325.98(66.65) | **<.001** | 9.99 | **<.001** | -123.6(65.61) | 0.158 | 180.31(55.38) | **0.004** | 303.91(71.39) | **<.001** |
| Posterior Cingulate | 29.78 | **<.001** | -281.36(72.49) | **<.001** | 312.88(61.18) | **<.001** | 594.25(78.88) | **<.001** | 28.72 | **<.001** | -295.16(71.78) | **<.001** | 287.8(60.59) | **<.001** | 582.96(78.11) | **<.001** |
| Frontal Pole | 13.43 | **<.001** | -61.92(23.29) | **0.028** | 65.98(19.66) | **0.003** | 127.9(25.35) | **<.001** | 10.67 | **<.001** | -57.6(27.75) | 0.109 | 75.53(23.42) | **0.004** | 133.13(30.2) | **<.001** |
| Parahippocampal | 20.35 | **<.001** | -179.9(44.09) | **<.001** | 127.06(37.22) | **0.002** | 306.96(47.98) | **<.001** | 28.21 | **<.001** | -138.02(34.36) | **<.001** | 138.66(29) | **<.001** | 276.68(37.39) | **<.001** |

| | | | | | | | | | | | | | | | | |
|---|---|---|---|---|---|---|---|---|---|---|---|---|---|---|---|---|
| Pars Orbitalis | 25.45 | **<.001** | -188.33(45.33) | **<.001** | 161.03(38.26) | **<.001** | 349.36(49.32) | **<.001** | 25.52 | **<.001** | -184.17(55.16) | **0.004** | 229.66(46.55) | **<.001** | 413.83(60.02) | **<.001** |
| Postcentral | 42.66 | **<.001** | -969.96(197.26) | **<.001** | 980.22(166.5) | **<.001** | 1950.18(214.65) | **<.001** | 53.09 | **<.001** | -897.23(188.73) | **<.001** | 1145.41(159.3) | **<.001** | 2042.64(205.37) | **<.001** |
| Pars Opercularis | 14.42 | **<.001** | -284.41(122.14) | **0.063** | 394.18(103.09) | **<.001** | 678.59(132.9) | **<.001** | 9.24 | **<.001** | -203.67(99.76) | 0.116 | 251.81(84.2) | **0.009** | 455.47(108.56) | **<.001** |
| Precuneus | 33.76 | **<.001** | -609.37(188.52) | **0.006** | 977.94(159.12) | **<.001** | 1587.31(205.14) | **<.001** | 31.79 | **<.001** | -705.26(193.03) | **0.002** | 911.16(162.92) | **<.001** | 1616.42(210.04) | **<.001** |
| Precentral | 36.28 | **<.001** | -881.89(193.57) | **<.001** | 894.38(163.38) | **<.001** | 1776.27(210.63) | **<.001** | 55.65 | **<.001** | -1147.23(198.6) | **<.001** | 1107.93(167.62) | **<.001** | 2255.16(216.1) | **<.001** |
| Pars Triangularis | 14.55 | **<.001** | -122.09(96) | 0.427 | 368.42(81.03) | **<.001** | 490.5(104.47) | **<.001** | 13.75 | **<.001** | -98.84(113.28) | 0.666 | 446.52(95.61) | **<.001** | 545.37(123.26) | **<.001** |
| Temporal Pole | 20.11 | **<.001** | -103.05(47.59) | 0.091 | 200.92(40.17) | **<.001** | 303.98(51.79) | **<.001** | 11.57 | **<.001** | -113.85(47.45) | 0.053 | 130.1(40.05) | **0.004** | 243.96(51.63) | **<.001** |
| Rostral Anterior Cingulate | 32.65 | **<.001** | -273.02(72.5) | **0.001** | 342.74(61.19) | **<.001** | 615.76(78.89) | **<.001** | 26.84 | **<.001** | -270.76(60.8) | **<.001** | 210.38(51.31) | **<.001** | 481.14(66.15) | **<.001** |
| Rostral Middle Frontal | 44.1 | **<.001** | -831.15(293.57) | **0.018** | 1880.84(247.78) | **<.001** | 2711.99(319.45) | **<.001** | 28.83 | **<.001** | -1287.69(317.96) | **<.001** | 1298.13(268.37) | **<.001** | 2585.82(345.99) | **<.001** |
| Superior Frontal | 43.82 | **<.001** | -2017.47(363.49) | **<.001** | 1671.94(306.8) | **<.001** | 3689.42(395.53) | **<.001** | 49.14 | **<.001** | -1936.87(362.04) | **<.001** | 1918.55(305.57) | **<.001** | 3855.42(393.96) | **<.001** |
| Supramarginal | 17.01 | **<.001** | -569.42(321.31) | 0.189 | 1288.49(271.2) | **<.001** | 1857.91(349.63) | **<.001** | 29.31 | **<.001** | -586.67(236.25) | **0.044** | 1223.98(199.41) | **<.001** | 1810.65(257.08) | **<.001** |
| Superior Parietal | 22.06 | **<.001** | -673.96(281.11) | 0.053 | 1214.25(237.27) | **<.001** | 1888.21(305.89) | **<.001** | 18.88 | **<.001** | -486.49(264.67) | 0.168 | 1109.88(223.39) | **<.001** | 1596.38(288) | **<.001** |
| Superior Temporal | 34.96 | **<.001** | -1032.71(208.32) | **<.001** | 856.32(175.83) | **<.001** | 1889.03(226.68) | **<.001** | 55.91 | **<.001** | -866.23(176.35) | **<.001** | 1097.47(148.84) | **<.001** | 1963.7(191.89) | **<.001** |
| Transverse Temporal | 21.27 | **<.001** | -120.12(31.05) | **<.001** | 98.57(26.21) | **<.001** | 218.69(33.79) | **<.001** | 29.85 | **<.001** | -91.74(22.78) | **<.001** | 95.8(19.23) | **<.001** | 187.54(24.79) | **<.001** |

**eTable 2.** Comparisons across subtypes of regional GM volumes at 2-year follow up

| | Omnibus Anova | | TD - S1 | | TD - S2 | | S1 - S2 | | Omnibus Anova | | TD - S1 | | TD - S2 | | S1 - S2 | |
|---|---|---|---|---|---|---|---|---|---|---|---|---|---|---|---|---|
| Region | F | p | Estimate (SE) | p | Estimate (SE) | p | Estimate (SE) | p | F | p | Estimate (SE) | p | Estimate (SE) | p | Estimate (SE) | p |
| | **Left Hemisphere** | | | | | | | | **Right Hemisphere** | | | | | | | |
| Nucleus Acumbens | 29.45 | **<.001** | -49.81(14.16) | **0.006** | 59.99(11.81) | <.001 | 109.8(14.95) | <.001 | 33.54 | <.001 | -31.16(13.22) | 0.072 | 71.5(11.03) | <.001 | 102.65(13.95) | <.001 |
| Amygdala | 19.7 | **<.001** | -67.67(31.58) | 0.106 | 125.26(26.34) | <.001 | 192.93(33.33) | <.001 | 24.28 | <.001 | -78.24(30.96) | 0.054 | 134.23(25.82) | <.001 | 212.48(32.68) | <.001 |
| Caudate | 25.05 | **<.001** | -245.35(69.4) | **0.006** | 259.14(57.89) | <.001 | 504.49(73.25) | <.001 | 24.52 | <.001 | -289.55(68.47) | 0.002 | 210.13(57.11) | 0.001 | 499.68(72.27) | <.001 |
| Hippocampus | 29.17 | **<.001** | -187.78(54.6) | **0.007** | 236.67(45.54) | <.001 | 424.45(57.63) | <.001 | 31.72 | <.001 | -192.95(55.39) | 0.006 | 252.79(46.2) | <.001 | 445.74(58.46) | <.001 |
| Pallidum | 12.56 | **<.001** | -85.27(34.28) | 0.057 | 91.42(28.59) | 0.005 | 176.69(36.18) | <.001 | 33 | <.001 | -110.82(29.04) | 0.004 | 130.75(24.22) | <.001 | 241.57(30.65) | <.001 |
| Putamen | 24.42 | **<.001** | -307.2(80.88) | **0.004** | 271.18(67.46) | <.001 | 578.37(85.37) | <.001 | 24.82 | <.001 | -304.3(81.01) | 0.004 | 277.93(67.57) | <.001 | 582.23(85.51) | <.001 |
| Thalamus | 45.8 | **<.001** | -394.94(105.78) | **0.004** | 614.25(88.23) | <.001 | 1009.19(111.65) | <.001 | 55.96 | <.001 | -274.26(91) | 0.019 | 643.63(75.91) | <.001 | 917.89(96.05) | <.001 |
| Ventral Diencephalon | 45.32 | **<.001** | -167.96(55.35) | **0.018** | 344.59(46.16) | <.001 | 512.55(58.42) | <.001 | 39.16 | <.001 | -220.24(54.92) | 0.003 | 276.36(45.81) | <.001 | 496.6(57.97) | <.001 |
| Banks of the Superior Temporal Sulcus | 14.87 | **<.001** | -237.89(81.6) | **0.022** | 216.96(68.06) | 0.005 | 454.85(86.12) | <.001 | 16.15 | <.001 | -224.91(70.85) | 0.014 | 185.96(59.1) | 0.006 | 410.87(74.79) | <.001 |
| Caudal Anterior Cingulate | 10.78 | **<.001** | -201.2(91.89) | 0.098 | 231.27(76.65) | 0.009 | 432.47(97) | <.001 | 6.1 | 0.003 | -209.82(82.68) | 0.054 | 76.43(68.97) | 0.51 | 286.26(87.27) | 0.003 |
| Caudal Middle Frontal | 22.22 | **<.001** | -911.8(227.97) | **0.003** | 649.41(190.15) | 0.003 | 1561.22(240.63) | <.001 | 26.52 | <.001 | -778.47(218.99) | 0.006 | 832.42(182.66) | <.001 | 1610.89(231.15) | <.001 |
| Cuneus | 8.32 | **<.001** | -239.2(97.67) | 0.059 | 169.35(81.47) | 0.098 | 408.55(103.09) | <.001 | 11.63 | <.001 | -164.68(106.71) | 0.305 | 328.76(89.01) | 0.001 | 493.44(112.64) | <.001 |
| Entorhinal | 3.14 | **0.045** | -28.91(61.08) | 0.884 | 108.57(50.94) | 0.089 | 137.48(64.46) | 0.086 | 2.14 | 0.12 | -46.34(57.09) | 0.704 | 71.18(47.62) | 0.299 | 117.52(60.26) | 0.127 |
| Fusiform | 12.45 | **<.001** | -438.38(208.93) | 0.114 | 588.77(174.27) | 0.003 | 1027.15(220.53) | <.001 | 15.34 | <.001 | -352.6(207.62) | 0.236 | 743.55(173.17) | <.001 | 1096.15(219.14) | <.001 |
| Isthmus Cingulate | 19.45 | **<.001** | -197.79(78.79) | 0.055 | 288.16(65.72) | <.001 | 485.95(83.16) | <.001 | 19.07 | <.001 | -162.93(73.31) | 0.094 | 276.57(61.15) | <.001 | 439.5(77.38) | <.001 |
| Insula | 45.55 | **<.001** | -308.72(113.72) | **0.036** | 721.39(94.85) | <.001 | 1030.11(120.03) | <.001 | 32.24 | <.001 | -441.39(116.36) | 0.004 | 509.03(97.06) | <.001 | 950.42(122.82) | <.001 |
| Inferior Parietal | 19.71 | **<.001** | -821.72(331.94) | 0.057 | 1223.95(276.87) | <.001 | 2045.68(350.36) | <.001 | 19.41 | <.001 | -1214.33(387.25) | 0.015 | 1211.77(323) | <.001 | 2426.1(408.74) | <.001 |
| Inferior Temporal | 22 | **<.001** | -820.8(283.33) | **0.023** | 1072.95(236.32) | <.001 | 1893.75(299.05) | <.001 | 20.47 | <.001 | -760.32(281.09) | 0.036 | 1052.58(234.45) | <.001 | 1812.89(296.69) | <.001 |
| Lingual | 13.98 | **<.001** | -440.14(182.65) | 0.064 | 518.73(152.35) | 0.003 | 958.87(192.79) | <.001 | 19.75 | <.001 | -564.8(188.62) | 0.019 | 631.48(157.33) | <.001 | 1196.27(199.09) | <.001 |
| Lateral Orbitofrontal | 38.19 | **<.001** | -367.02(140.36) | **0.044** | 800.76(117.07) | <.001 | 1167.78(148.15) | <.001 | 35.46 | <.001 | -453.4(138.69) | 0.01 | 699.21(115.68) | <.001 | 1152.61(146.39) | <.001 |
| Lateral Occipital | 17.47 | **<.001** | -591.25(270.96) | 0.098 | 966.97(226.01) | <.001 | 1558.22(286) | <.001 | 16.59 | <.001 | -645.68(302.57) | 0.106 | 1056.28(252.37) | <.001 | 1701.97(319.36) | <.001 |
| Medial Orbitofrontal | 15.2 | **<.001** | -332.24(108.84) | **0.018** | 276.17(90.79) | 0.008 | 608.41(114.89) | <.001 | 24.91 | <.001 | -234.87(103.03) | 0.084 | 462.91(85.94) | <.001 | 697.78(108.75) | <.001 |
| Middle Temporal | 40.05 | **<.001** | -996.71(263.06) | **0.004** | 1358.15(219.42) | <.001 | 2354.86(277.66) | <.001 | 44.26 | <.001 | -1188.56(259.55) | 0.001 | 1291.97(216.49) | <.001 | 2480.52(273.95) | <.001 |
| Paracentral | 17.89 | **<.001** | -264.49(89.54) | **0.021** | 278.07(74.69) | 0.001 | 542.56(94.51) | <.001 | 21.44 | <.001 | -316.36(94.8) | 0.009 | 315.94(79.07) | <.001 | 632.31(100.06) | <.001 |
| Pericalcarine | 7.51 | **<.001** | -131.29(76) | 0.228 | 165.07(63.39) | 0.029 | 296.36(80.22) | <.001 | 8.04 | <.001 | -73.8(78.58) | 0.647 | 217.38(65.55) | 0.004 | 291.18(82.94) | 0.002 |
| Posterior Cingulate | 19.77 | **<.001** | -264.38(88.21) | **0.019** | 296.06(73.57) | <.001 | 560.44(93.1) | <.001 | 22.65 | <.001 | -301.46(86.91) | 0.006 | 292.11(72.49) | <.001 | 593.57(91.73) | <.001 |
| Frontal Pole | 6.22 | **0.002** | -44.33(29.78) | 0.329 | 59.5(24.84) | 0.049 | 103.83(31.44) | 0.003 | 12.64 | <.001 | -72.44(34.8) | 0.116 | 101.3(29.02) | 0.002 | 173.75(36.73) | <.001 |
| Parahippocampal | 14.38 | **<.001** | -151.51(51.78) | **0.022** | 139.1(43.19) | 0.005 | 290.61(54.65) | <.001 | 13.88 | <.001 | -137.66(41.78) | 0.01 | 92.77(34.85) | 0.025 | 230.43(44.1) | <.001 |
| Pars Orbitalis | 26.79 | **<.001** | -181.56(52.05) | **0.006** | 204.54(43.42) | <.001 | 386.1(54.94) | <.001 | 13.54 | <.001 | -135.5(73.32) | 0.185 | 232.96(61.15) | <.001 | 368.46(77.39) | <.001 |

| | | | | | | | | | | | | | | | | |
|---|---|---|---|---|---|---|---|---|---|---|---|---|---|---|---|---|
| Postcentral | 29.35 | **<.001** | -745.13(240.99) | **0.016** | 1087.64(201) | <.001 | 1832.77(254.36) | <.001 | 30.79 | <.001 | -887.8(240.09) | 0.004 | 1013.61(200.26) | <.001 | 1901.4(253.42) | <.001 |
| Pars Opercularis | 10.78 | **<.001** | -382.09(155.1) | 0.058 | 344(129.37) | 0.025 | 726.1(163.71) | <.001 | 10.27 | <.001 | -334.56(122.94) | 0.036 | 245.3(102.54) | 0.049 | 579.87(129.76) | <.001 |
| Precuneus | 24.09 | **<.001** | -508.63(223.38) | 0.084 | 976.89(186.32) | <.001 | 1485.52(235.78) | <.001 | 23.61 | <.001 | -619.42(229.87) | 0.036 | 928.97(191.73) | <.001 | 1548.4(242.63) | <.001 |
| Precentral | 23.33 | **<.001** | -690.39(243.66) | **0.027** | 969.76(203.24) | <.001 | 1660.15(257.19) | <.001 | 33.05 | <.001 | -1010.63(254.24) | 0.003 | 1093.25(212.06) | <.001 | 2103.89(268.35) | <.001 |
| Pars Triangularis | 14.64 | **<.001** | -95.91(109.62) | 0.673 | 423.92(91.43) | <.001 | 519.83(115.7) | <.001 | 10.85 | <.001 | -125.19(139.84) | 0.668 | 464.51(116.64) | <.001 | 589.69(147.6) | <.001 |
| Temporal Pole | 6.73 | **0.001** | -90.56(63.57) | 0.355 | 138.88(53.02) | 0.028 | 229.44(67.1) | 0.002 | 6.79 | 0.001 | -107.8(55.7) | 0.157 | 105.98(46.46) | 0.063 | 213.77(58.79) | 0.001 |
| Rostral Anterior Cingulate | 24.91 | **<.001** | -333.31(88.96) | **0.004** | 309.32(74.2) | <.001 | 642.63(93.9) | <.001 | 22.14 | <.001 | -296.16(72.6) | 0.003 | 199.94(60.55) | 0.004 | 496.1(76.63) | <.001 |
| Rostral Middle Frontal | 32.94 | **<.001** | -853.7(364.36) | 0.074 | 1935.25(303.91) | <.001 | 2788.95(384.58) | <.001 | 22.27 | <.001 | -1008.49(403.18) | 0.056 | 1614.89(336.29) | <.001 | 2623.38(425.55) | <.001 |
| Superior Frontal | 37.86 | **<.001** | -2010.34(443.12) | **0.001** | 1919.29(369.6) | <.001 | 3929.63(467.71) | <.001 | 38.73 | <.001 | -1941.08(448.61) | 0.002 | 2090.22(374.18) | <.001 | 4031.31(473.51) | <.001 |
| Supramarginal | 15.56 | **<.001** | -633.39(367.27) | 0.228 | 1331.16(306.34) | <.001 | 1964.54(387.66) | <.001 | 23.11 | <.001 | -602.04(275.68) | 0.098 | 1204.98(229.94) | <.001 | 1807.02(290.98) | <.001 |
| Superior Parietal | 16.74 | **<.001** | -489.77(343.12) | 0.355 | 1339.52(286.19) | <.001 | 1829.29(362.16) | <.001 | 8.69 | <.001 | -392.22(324.32) | 0.477 | 874.39(270.52) | 0.005 | 1266.6(342.32) | <.001 |
| Superior Temporal | 25.2 | **<.001** | -968(255.21) | **0.004** | 895.97(212.87) | <.001 | 1863.97(269.37) | <.001 | 41.05 | <.001 | -744.21(217.35) | 0.007 | 1202.33(181.29) | <.001 | 1946.54(229.41) | <.001 |
| Transverse Temporal | 14.34 | **<.001** | -109.93(39.49) | **0.031** | 106.81(32.93) | 0.005 | 216.74(41.68) | <.001 | 24 | <.001 | -90.37(28.83) | 0.015 | 112.63(24.05) | <.001 | 203(30.43) | <.001 |

**eFigure 1.** Longitudinal ComBat Harmonization

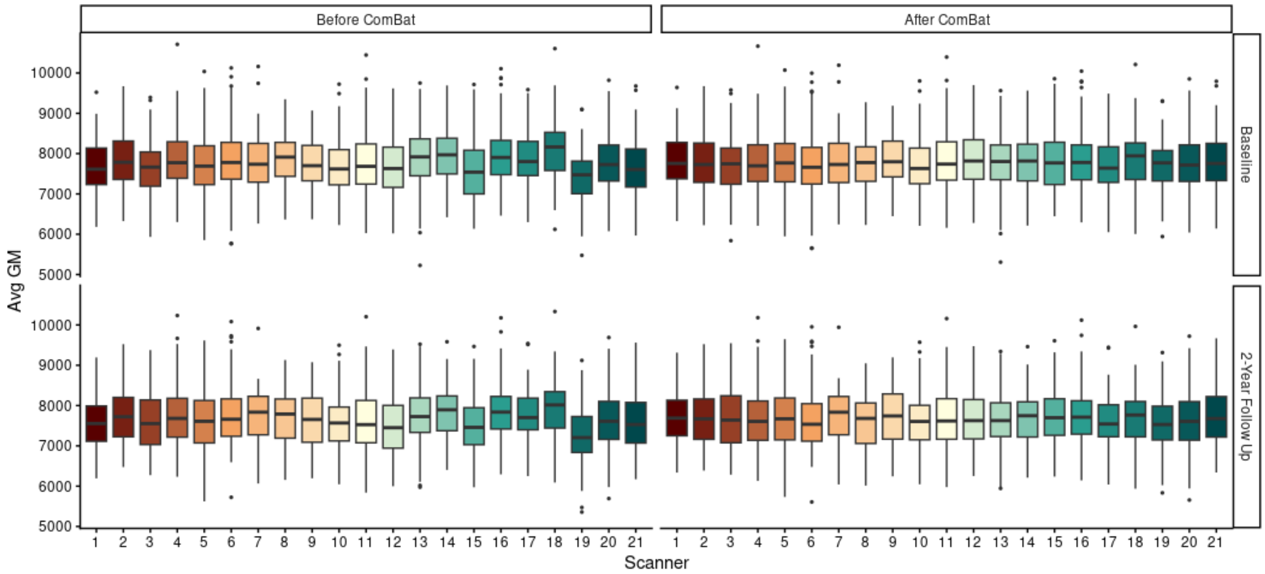

**Note.** Longitudinal ComBat harmonization was applied to account for site and scanner effects in imaging data. We implemented Longitudinal ComBat on all ROI's for both cortical and subcortical GM, leveraging a generalized additive mixed effect modeling strategy to allow for nonlinear effects of age.

**eFigure 2.** A schematic outline of the HYDRA algorithm.

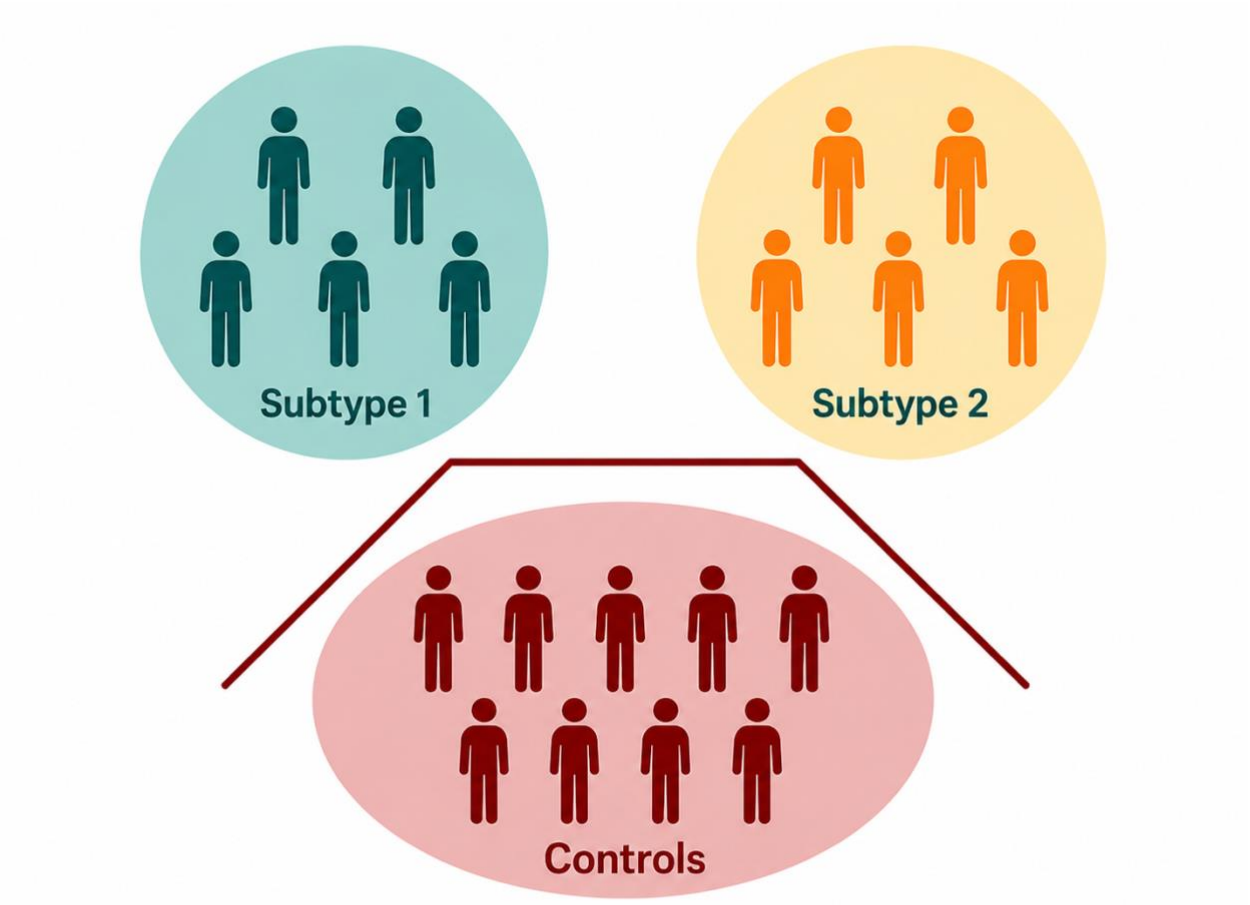

**Note.** The polytope (represented by the red line) separates the typically developing controls (in red) from each of the subtypes using 2 separate Support Vector Machine hyperplanes.

**eFigure 3.** Bi-variate correlations between study variables used for subtype validation.

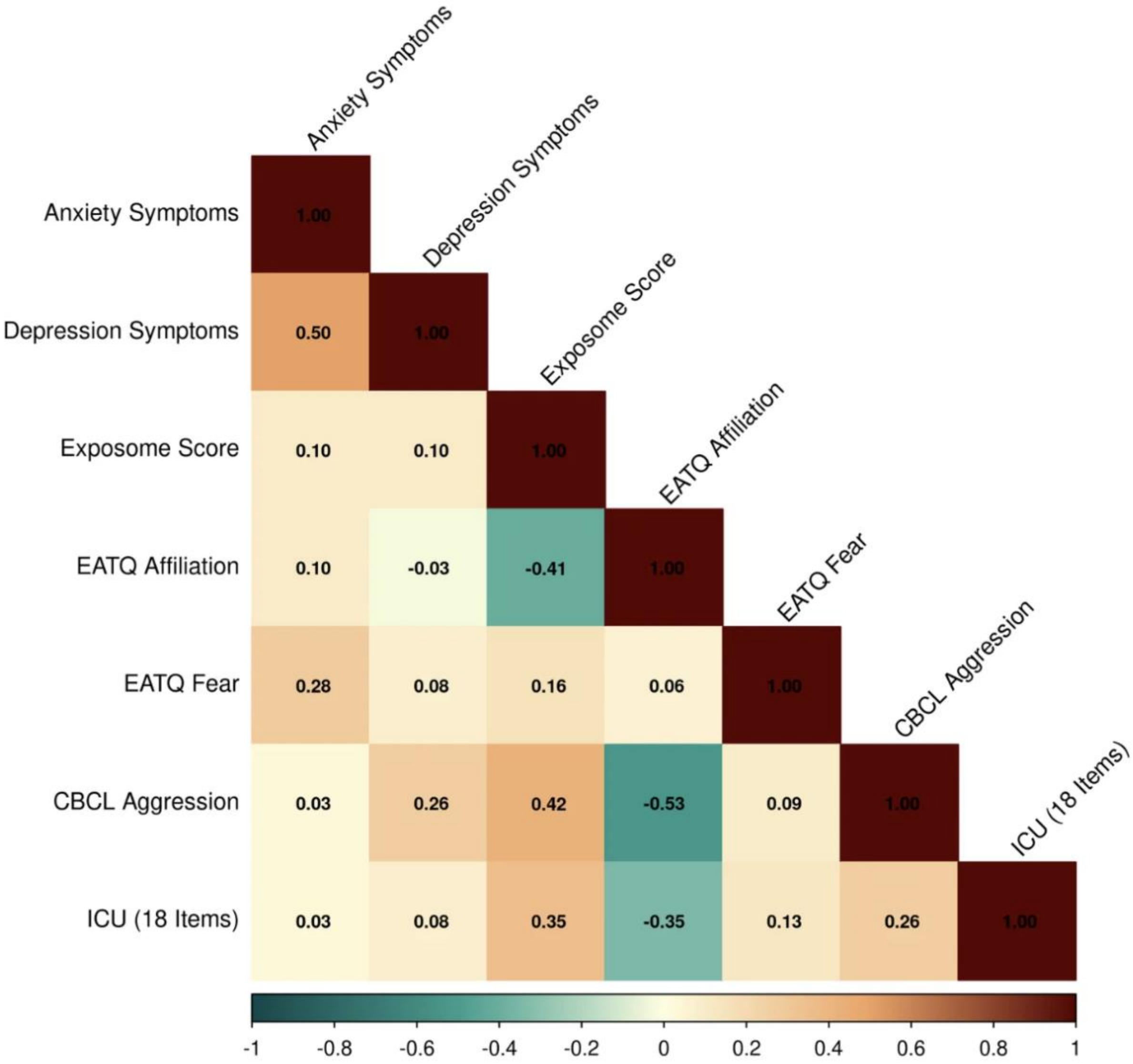

**eFigure 4.** Subtype differences on the 18-Item Inventory of Callous Unemotional Traits

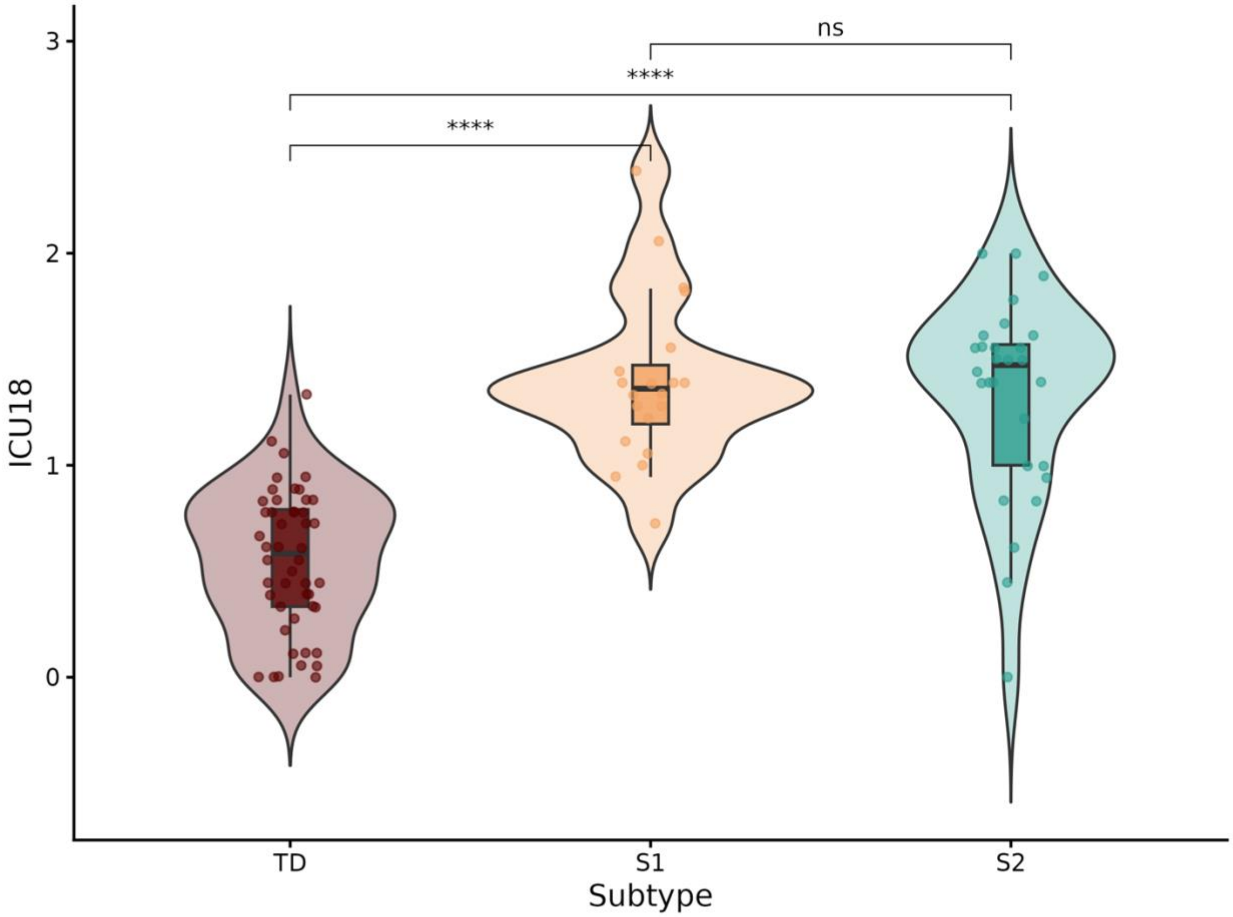

**Note.** ANCOVA models revealed that subtypes differed on levels of CU traits as measured by 18 items from the Inventory of Callous Unemotional Traits ($F(2,91) = 50.6218$, $p<.001$). Both S1 and S2 subtypes evidenced higher levels of CU traits than TD, but did not differ significantly from each other.

**eFigure 5.** Profile stability across T1 subtypes.

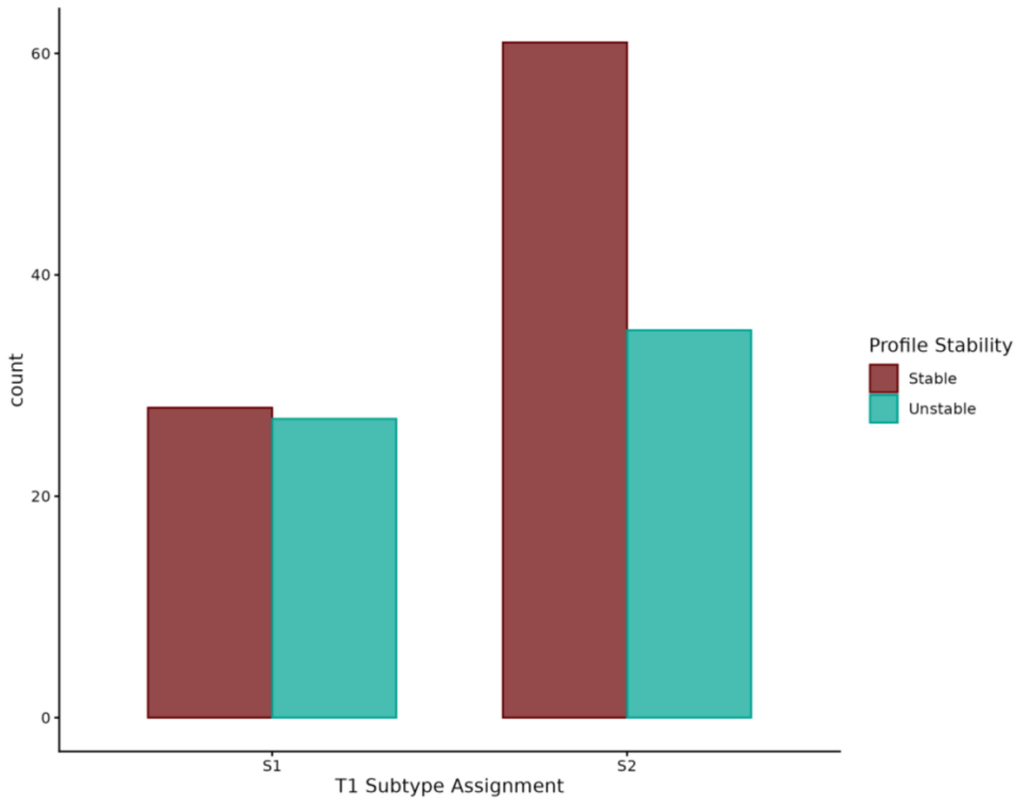

**Note.** A chi-square test of independence indicated a significant association between the variables in subtype assignment at T1 and subtype stability at T2, $\chi^2(3, N = 151) = 67.76$, $p < .001$, such that participants in S1 were more likely to be assigned to a "both" or "neither" profile at T2 than participants in S2.

**Supplemental References**

Frick, P. J. (2004). Inventory of Callous–Unemotional Traits. https://doi.org/10.1037/t62639-000

Hoffman, E. A., Clark, D. B., Orendain, N., Hudziak, J., Squeglia, L. M., & Dowling, G. J. (2019). Stress exposures, neurodevelopment and health measures in the ABCD study. Neurobiology of Stress, 10, 100157. https://doi.org/10.1016/j.ynstr.2019.100157

Shmilovici, A. (2005). Support Vector Machines. In O. Maimon & L. Rokach (Eds.), Data Mining and Knowledge Discovery Handbook (pp. 257–276). Springer US. https://doi.org/10.1007/0-387-25465-X_12

Wen, J., Fu, C. H. Y., Tosun, D., Veturi, Y., Yang, Z., Abdulkadir, A., Mamourian, E., Srinivasan, D., Skampardoni, I., Singh, A., Nawani, H., Bao, J., Erus, G., Shou, H., Habes, M., Doshi, J., Varol, E., Mackin, R. S., Sotiras, A., … iSTAGING consortium, A., BIOCARD, and BLSA. (2022). Characterizing Heterogeneity in Neuroimaging, Cognition, Clinical Symptoms, and Genetics Among Patients With Late-Life Depression. JAMA Psychiatry, 79(5), 464–474. https://doi.org/10.1001/jamapsychiatry.2022.0020
